# *ACTN3* genotype influences androgen response in skeletal muscle

**DOI:** 10.1101/2024.04.25.591034

**Authors:** Kelly N. Roeszler, Michael See, Lyra R. Meehan, Giscard Lima, Alexander Kolliari-Turner, Sarah E. Alexander, Shanie Landen, Harrison D. Wood, Chrystal F. Tiong, Weiyi Chen, Tomris Mustafa, Peter J. Houweling, Nir Eynon, Severine Lamon, Yannis Pitsiladis, David J. Handelsman, Fernando J. Rossello, Mirana Ramialison, Kathryn N. North, Jane T. Seto

## Abstract

Androgens are vital for the maintenance of muscle mass and their anabolic effects are primarily exerted through the androgen receptor (AR). Accumulating evidence in humans and mice suggests that circulating androgens, AR and androgen response are influenced by *ACTN3 (*α- actinin-3), also known as “the gene for speed”. One in 5 people worldwide are α-actinin-3 deficient due to homozygous inheritance of a common null polymorphism (577X) in *ACTN3*. In this study, we show that α-actinin-3 deficiency decreases baseline AR in skeletal muscles of mice and humans, in both males and females, and that AR expression directly correlates with *ACTN3* in a dosage dependent manner. We further demonstrate in *Actn3* knockout mice that α- actinin-3 deficiency increases muscle wasting induced by androgen deprivation and reduces the muscle hypertrophic response to dihydrotestosterone and this is mediated by differential activation of pathways regulating amino acid metabolism, intracellular transport, MAPK signalling, autophagy, mitochondrial activity and calcineurin signalling. Gene set enrichment and protein analyses indicate that the absence of α-actinin-3 results in a failure to coactivate many of these pathways in response to changes in androgens, and relies on leveraging mitochondrial remodelling and calcineurin signalling to restore muscle homeostasis. We further identified 7 genes that are androgen sensitive and α-actinin-3-dependent in expression, and whose functions correspond to these processes. Our results highlight the pivotal role of α- actinin-3 in various processes associated with the regulation of protein turnover and muscle mass, and suggest that *ACTN3* genotype is a genetic modifier of androgen action in skeletal muscle.

## Introduction

Androgens orchestrate the development and maintenance of masculine reproductive characteristics and have well characterised anabolic actions in skeletal muscle, heart, and bone^1^. Testosterone and its more potent metabolite dihydrotestosterone (DHT) are the primary androgens in male mammals and are present at low levels in females. Testosterone decline (due to hypogonadism, ageing or androgen deprivation therapy to treat prostate cancer) is associated with muscle wasting^2,3^, while testosterone replacement therapy is prescribed at physiological doses to men whose endogenous testosterone production is severely limited to maintain or restore muscle mass^1^.

The anabolic actions of androgens in muscles are in part exerted by binding to the androgen receptor (AR) as a ligand-activated transcription factor activating androgen action. Androgen action involves stimulation of protein synthesis pathways (such as mTORC1 and calcineurin signalling), and inhibition of protein degradation (via suppressing E3-ubiquitin ligase genes *Fbxo32* and *Trim63*)^3,4^. Androgens also directly promote expression of target genes such as myostatin (*Mstn*), a key repressor of muscle growth as a negative feedback mechanism to restrain unlimited muscle growth^5^, as well as genes involved in induction of autophagy and myoblast proliferation^6–8^. In addition, activation of AR involves recruitment of coactivators such as α-actinin-2 and glucocorticoid receptor interacting protein 1 (GRIP1) to facilitate transcription of target genes^9^.

α-Actinin-2 (*ACTN2*) and α-actinin-3 (*ACTN3*) are major components of the muscle contractile apparatus. α-Actinin-2 is present in all muscle fibres, while α-actinin-3 has developed specialised expression in only type 2 (fast-twitch, glycolytic) fibres, which are important for rapid, repetitive muscle contractions and for activities that require speed and power^10^. A common null polymorphism in *ACTN3* (R577X) arose in modern humans and underwent positive selection ∼10-15,000 years ago^11,12^. One in 5 people worldwide are completely α-actinin-3 deficient due to inheriting two copies of the null variant (577X) in the *ACTN3* gene^13^. α-Actinin-3 deficiency does not cause disease because of compensatory upregulation of α-actinin-2, which shares high sequence similarity^10^. However, the absence of α-actinin-3 is detrimental to sprint and power performance in elite athletes and the general population^14–17^. Among the elderly, α-actinin-3 deficiency is associated with significantly lower strength and higher frailty scores in an elderly Chinese population^18^, and a meta-analysis using two large independent cohorts of Caucasian post-menopausal women showed that carriage of the *ACTN3* 577X allele increases the risk of falling by 33%^19^.

Accumulating evidence in humans and mouse models suggest a relationship between α-actinin- 3 expression and androgen response. In a cohort of elite male and female Russian athletes, carriage of the *ACTN3* 577R allele was associated with significantly higher testosterone levels compared to XX individuals^20^. Testicular feminised mice (*Tfm;* which have an inactivating mutation in AR) and hypogonadal mice (*hpg*; which lack gonadotropin and sex steroid production) are both androgen deficient and both show significantly reduced *Actn3* expression in the testis, while testosterone treatment of *hpg* mice upregulated *Actn3,* confirming that *Actn3* is an androgen-regulated target gene in the testis^21^. Comparison of AR knockout (ARKO) mice with the *Actn3* KO mouse model also revealed many similarities in muscle characteristics. Relative to wild-type (WT) mice, both male ARKO and *Actn3* KO mice demonstrate decreased body weight, reductions in hind-limb muscle mass and strength, increased calcineurin signalling, increased expression of slow-twitch contractile proteins and enhanced fatigue resistance^22–24^. Similarly, both male *Actn3* KO and the fast-twitch muscle specific ARKO (fmARKO) show reduced bone mineral density and reduced baseline expression of *Mstn, Fbxo32* and *Trim63,* critical genes that regulate and maintain muscle mass^25–27^. These results suggest that changes in androgen response with *ACTN3* genotype underlie the alterations in muscle performance associated with α-actinin-3 deficiency.

On this basis, we hypothesise that *ACTN3* genotype will alter baseline AR expression and consequently, the muscle wasting response to androgen deprivation and muscle growth response to androgen treatments. We have previously shown that α-actinin-3 deficiency changes how muscles adapt following denervation and immobilisation and protects against dexamethasone-induced muscle wasting through increases in protein synthesis pathways mTORC1 and calcineurin signalling and suppressing protein degradation^26,28^. In this study, we examined the effect of α-actinin-3 deficiency on AR signalling and the skeletal muscle response to changes in circulating androgens.

## Results

### AR protein expression is reduced with α-actinin-3 deficiency in human skeletal muscle

We performed a trans-eQTL scan in the GTEx skeletal muscle dataset (GTEx Analysis Release V8 (dbGaP Accession phs000424.v8.p2)) to determine if the *ACTN3* R577X variant (rs1815739) is associated with the expression of *ACTN3, ACTN2* and *AR* (Fig 1a). Consistent with our previous analysis using an earlier GTEx release^29^, the R577X variant is significantly associated with *ACTN3* gene expression in skeletal muscle (*P* = 7.1 × 10^−141^), but not for *ACTN2.* Normalised gene expression of *AR* is also not significantly different between *ACTN3* R577X genotypes (Fig 1a). However, qPCR analysis of *AR* in vastus lateralis muscles from a small cohort of moderately trained Caucasian men (aged 18-47; *N* = 24) and women (aged 21- 45; *N* = 20) show a trend for reduced *AR* expression in 577XX individuals compared to RR+RX (Supplementary Fig 1a). We then examined protein expression of AR in human skeletal muscles from a cohort of young Caucasian men (aged 22-42, *N =* 12) and women (aged 18-37, *N* = 24) to determine if AR levels are altered in association with *ACTN3* R577X genotype. Our results show that muscle AR is significantly reduced in XX individuals compared to RR in both male and female cohorts by 65% (*P* = 0.0043) and 72% (*P* = 0.0112), respectively (Fig 1b).

**Fig 1.**
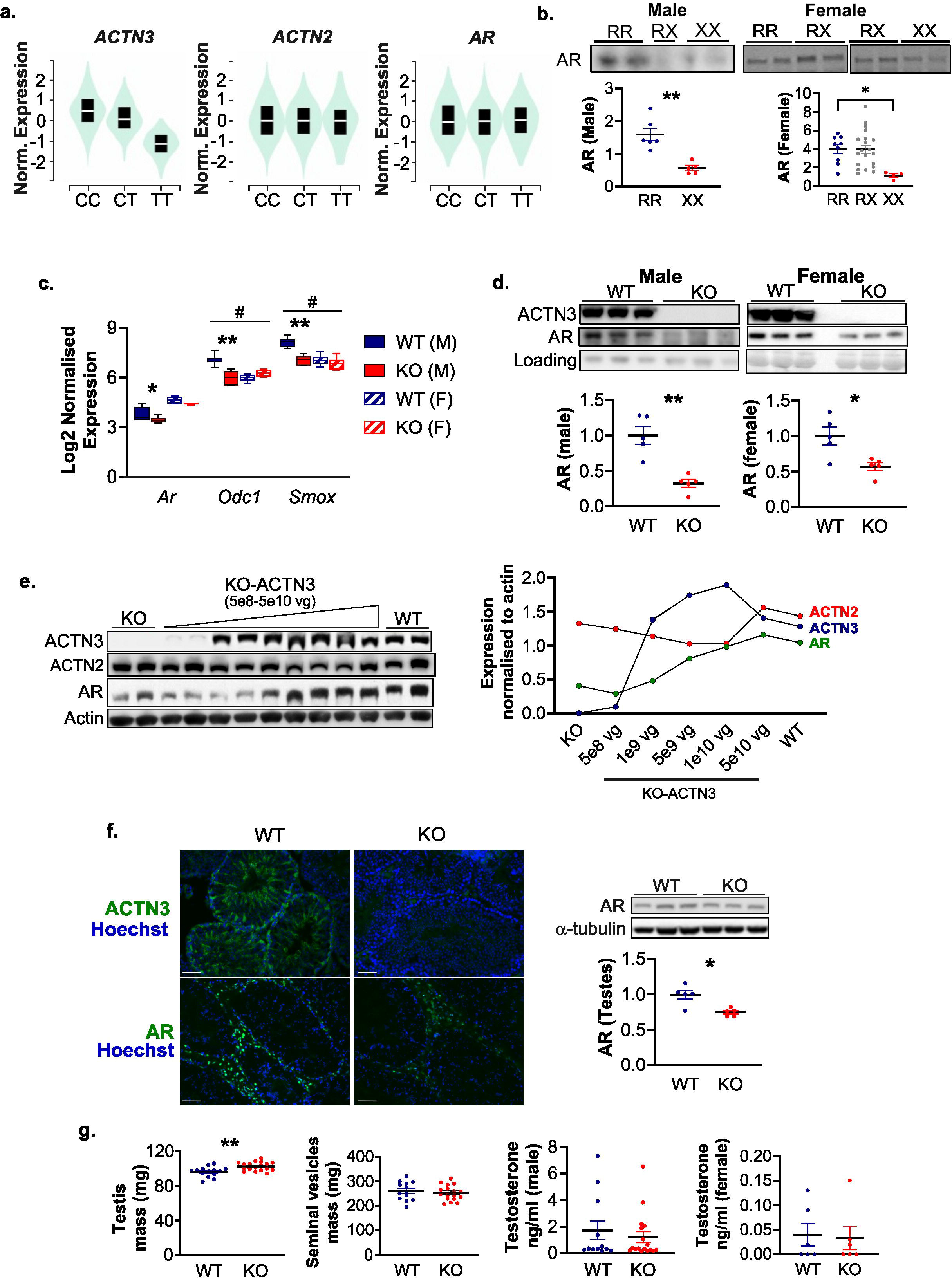
AR protein expression is reduced with α-actinin-3 deficiency in skeletal muscle and testis. a) Trans-eQTL scan in the GTEx skeletal muscle dataset show that the R577X variant is significantly associated with *ACTN3* gene expression but not for *ACTN2* or *AR.* b) Protein expression of AR is significantly reduced in muscles from 577XX individuals compared to 577RR in males and females. c) Expression of *Ar* and *Ar-*responsive genes (*Odc1, Smox*) is lower in *Actn3* KO muscles relative to WT in male but not female mice. d) AR protein expression is lower in gastrocnemius muscles from male and female *Actn3* KO mice compared to WT. e) Delivery of rAAV*-*CMV-*ACTN3* (5e8 – 5e10 vg) in *Actn3* KO muscles increased α- actinin-3 expression up to 1e10 vg and decreased α-actinin-2, while AR expression is positively correlated with vector dosage. f) Immunohistochemistry show absence of ACTN3 in KO testes and reduced AR staining compared to WT; reduced AR expression in KO testis is quantified by western blot. g) Testis mass is greater in KO mice compared to WT, but seminal vesicle mass and serum testosterone, as measured by radioimmunoassay assay (males) and mass spectrometry (females), are not different between *Actn3* genotypes. Data are represented as mean ± SEM. \**p* < 0.05, \*\**p* < 0.01 by Mann-Whitney U test; #*p* < 0.05 by two-way ANOVA (c).

### AR is reduced in Actn3 knockout mouse muscles and testes and is sensitive to increases in α- actinin-3 expression

We further examined the expression of *Ar* and *Ar-*responsive polyamine biosynthesis genes (*Odc1, Smox*)^6^ in male and female WT and *Actn3* KO mouse muscles at baseline. Expression of these genes are lower in *Actn3* KO muscles compared to WT in male but not female mice (Fig 1c); two-way ANOVA showed significant main and interaction effects between *Actn3* genotype and sex for *Odc1* (F(1, 21) = 31.76, *P* < 0.0001) and *Smox* (F(1, 21) = 12.55, *P* = 0.0019). At the protein level, *Actn3* KO gastrocnemius muscles showed 67.6% and 43% lower levels of AR in male (*P*= 0.0079) and female mice (*P*= 0.0317), respectively, compared to WT (Fig 1d). Consistent with reductions in total AR expression with α-actinin-3 deficiency, phosphorylation of AR at Ser^213/210^ and Ser^791/790^, which marks AR for degradation^30^, is higher in *Actn3* KO muscles by 59.1% (Supplementary Fig 1b).

Estrogen receptor (ER) and thyroid receptor (TR) signalling were also examined, given their involvement in skeletal muscle myogenesis, metabolism and contraction^31^, and α-actinin-2 and -4 have been shown to directly interact with these nuclear receptors and potentiate their activation^9,32^. In contrast to AR, expression of genes associated with ER (*Esr1, Esrra, Esrrb, Esrrg*) and TR (*Thra, Thrb, Thrsp*) are similar between WT and *Actn3* KO muscles, regardless of sex (Supplementary Table 1). ER-α and ER-β protein expression is also similar in WT and *Actn3* KO muscles (Supplementary Fig 1c, d).

To determine if AR protein expression is associated with α-actinin-3 dosage, variable doses of rAAV*-*CMV-*ACTN3* (5e8 – 5e10 vg) were delivered by intramuscular injection into the tibialis anterior muscles of *Actn3* KO mice, and AR expression assessed (Fig 1e). Results show a rise in α-actinin-3 expression in KO muscles with escalating doses of rAAV-CMV-*ACTN3* up to 1e10 vg, dropping at 5e10 vg. Consistent with the regulation of total sarcomeric α-actinin content^17^, a reciprocal decrease in α-actinin-2 expression is also observed up to 1e10 vg, which then increases at 5e10 vg. In contrast, AR expression is positively correlated with rAAV-CMV- *ACTN3* dosage.

We further examined the effects of α-actinin-3 deficiency on AR expression in the mouse testis since GTEX analysis indicates that *ACTN3* R577X also influences *ACTN3* expression in human testis (Supplementary Fig 1e). Immunostaining in WT testis shows α-actinin-3 expression within the seminiferous tubules, while AR labelling is strongest in the interstitial compartment that coincide with the location of Leydig cells^33^. In contrast, α-actinin-3 is absent in the testis of *Actn3* KO mice, while AR immunolabelling is reduced (Fig 1f). Furthermore, testis mass is 6.5% higher in *Actn3* KO mice compared to WT (*P*= 0.0032), but seminal vesicle mass is not different between WT or *Actn3* KO mice (Fig 1g). Despite changes in AR expression in the testis with α-actinin-3 deficiency, radioimmunoassay assay and mass spectrometry show similar levels of serum testosterone and luteinising hormone between *Actn3* genotypes in male and female mice (Fig 1g, Supplementary Fig 1f).

### α-Actinin-3 deficiency increases muscle wasting induced by androgen deprivation

To determine if baseline reductions in AR signalling with α-actinin-3 deficiency influence the skeletal muscle response to androgen deprivation, orchidectomy (ORX) was performed on adult male WT and *Actn3* KO mice (aged 8-10 weeks) and compared to sham operated mice. After 12 weeks, orchidectomy induced significant reductions in both WT-ORX and *Actn3* KO- ORX mice relative to sham controls in body mass, lean mass, bone mineral density (BMD), bone mineral content (BMC), the mass of seminal vesicles and the levator ani bulbocavernosus (LABC) muscle, as well as an increase in % fat mass, confirming complete androgen deprivation in both genotypes (Fig 2a-c, Supplementary Table 2).

**Fig 2.**
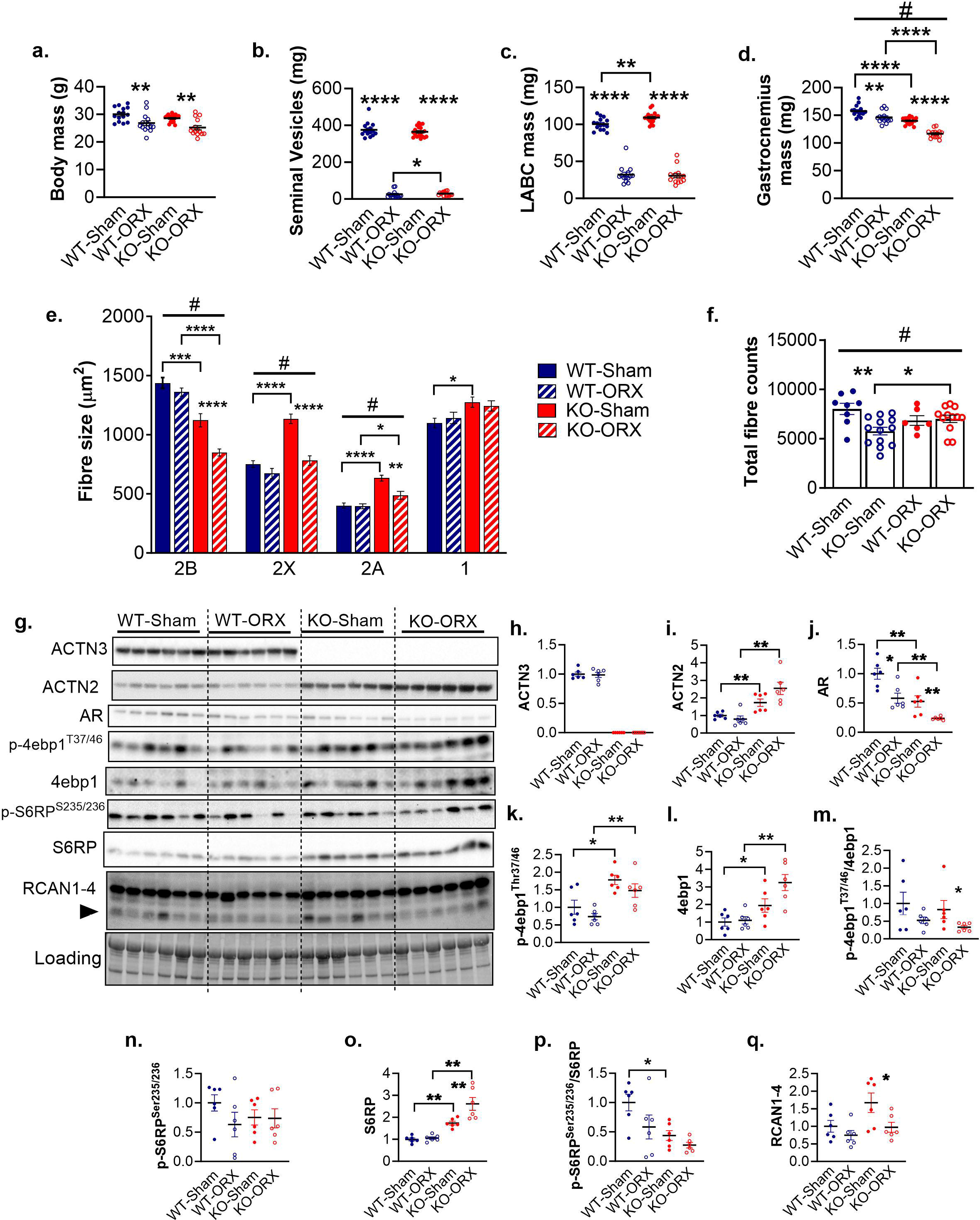
α-Actinin-3 deficiency differentially alters the muscle wasting and calcineurin signalling response induced by androgen deprivation. Orchidectomy decreased a) body mass, b) seminal vesicle, c) levator ani bulbocavernosus (LABC), and d) gastrocnemius mass of WT and *Actn3* KO mice; KO showed greater atrophy for the gastrocnemius muscle compared to WT. e) The size of fast 2B, 2X and 2A fibres is reduced in KO-ORX but not WT- ORX gastrocnemius muscles compared to controls. f) Muscles of orchidectomised WT but not KO show significant reductions in fibre count. g) Western blot analyses quantify the effect of orchidectomy in WT and KO gastrocnemius muscles on the expression of h) ACTN3, i) ACTN2, j) AR, k) p-4ebp1^Thr37/46^, l) total 4ebp1, m) ratio of p-4ebp1^Thr37/46^/4ebp1, n) p- S6RP^Ser235/236^, o) total S6RP, p) ratio of p-S6RP^Ser235/236^/S6RP, and q) RCAN1-4. Data are represented as mean ± SEM. \**p* < 0.05, \*\**p* < 0.01, \*\*\**p* < 0.001, \*\*\*\**p* < 0.0001 by Mann-Whitney U test; #*p* < 0.05 by two-way ANOVA (d-f).

Orchidectomy also induced significant decreases in the mass of all hindlimb muscles examined in both WT-ORX and KO-ORX, except for the soleus; there is also no effect on heart mass (Supplementary Table 3). However, KO-ORX mice showed significantly greater atrophy for the gastrocnemius muscle compared to WT-ORX (WT-ORX: -7.3%, KO-ORX: -16.7%, F(1, 59) = 7.666, *P* = 0.0075) (Fig 2d). Similar trends are observed in the quadriceps, tibialis anterior, extensor digitorum longus and spinalis, suggesting that α-actinin-3 deficiency increases muscle wasting induced by androgen deprivation.

### Actn3 KO-ORX muscle wasting is mediated by fibre atrophy and is associated with a concomitant decrease in calcineurin activity

Analysis of fibre size and number in the gastrocnemius muscle indicates differential mechanisms mediating muscle atrophy in WT-ORX and KO-ORX (Fig 2e-f). Two-way ANOVA confirmed a significant interaction between the effects of *Actn3* genotype and orchidectomy on the size of fast 2B (F(1, 47) = 5.239, *P* = 0.0266), 2X (F(1, 42) = 11.62, *P* = 0.0015) and 2A fibres (F(1, 40) = 6.561, *P* = 0.0143) with KO-ORX muscles showing significantly smaller fast 2B, 2X and 2A fibres compared to KO-Sham, while the size of fast and slow fibres is not altered in WT following orchidectomy (Fig 2e). In contrast, orchidectomy significantly decreased total fibre number in WT-ORX but not *Actn3* KO-ORX; two-way ANOVA showed significant interaction between *Actn3* genotype and effect of orchidectomy (F(1,35) = 6.687, *P* = 0.014) (Fig 2f). There is no change in fibre type proportions in either genotype (Supplementary Fig 2).

Further assessment of both WT-ORX and KO-ORX gastrocnemius muscles showed significant reductions in AR expression in both genotypes compared to sham controls (Fig 2g, i). Evaluation of the downstream changes in mTORC1 activity, which regulates cell size, metabolism and growth, showed similar reductions in the ratio of phosphorylated to total 4ebp1 and S6RP expression in both genotypes (Fig 2g-p). We also examined RCAN1-4 expression, a marker of calcineurin activity, since calcineurin mediates the androgen-induced hypertrophic response of myotubes^4^, and is also significantly increased in α-actinin-3 deficient muscles at baseline, and in response to immobilisation and denervation and exercise training^24,28^. There is a trend for a differential *Actn3* genotype effect on RCAN1-4 expression in response to androgen deprivation (F(1, 20) = 1.384, *P* = 0.2532); RCAN1-4 levels are similar between WT- ORX and WT-Sham but are significantly decreased in KO-ORX relative to KO-Sham (Fig 2q).

### α-Actinin-3 deficiency alters the transcriptional response to androgen deprivation

We further performed whole transcriptomic profiling to examine the mechanisms underlying the differential muscle wasting response with α-actinin-3 deficiency following orchidectomy (Fig 3). Principle component analysis of the filtered and processed count data confirmed the experimental grouping of gene expression by *Actn3* genotype and Sham/ORX treatment (Fig 3a). Differential expression testing for effects of orchidectomy identified 1113 genes in WT and 1116 genes in *Actn3* KO muscles that were differentially expressed (*q* ≤ 0.01) in the expression, with 492 of these genes common between *Actn3* genotypes (Fig 3b).

**Fig 3.**
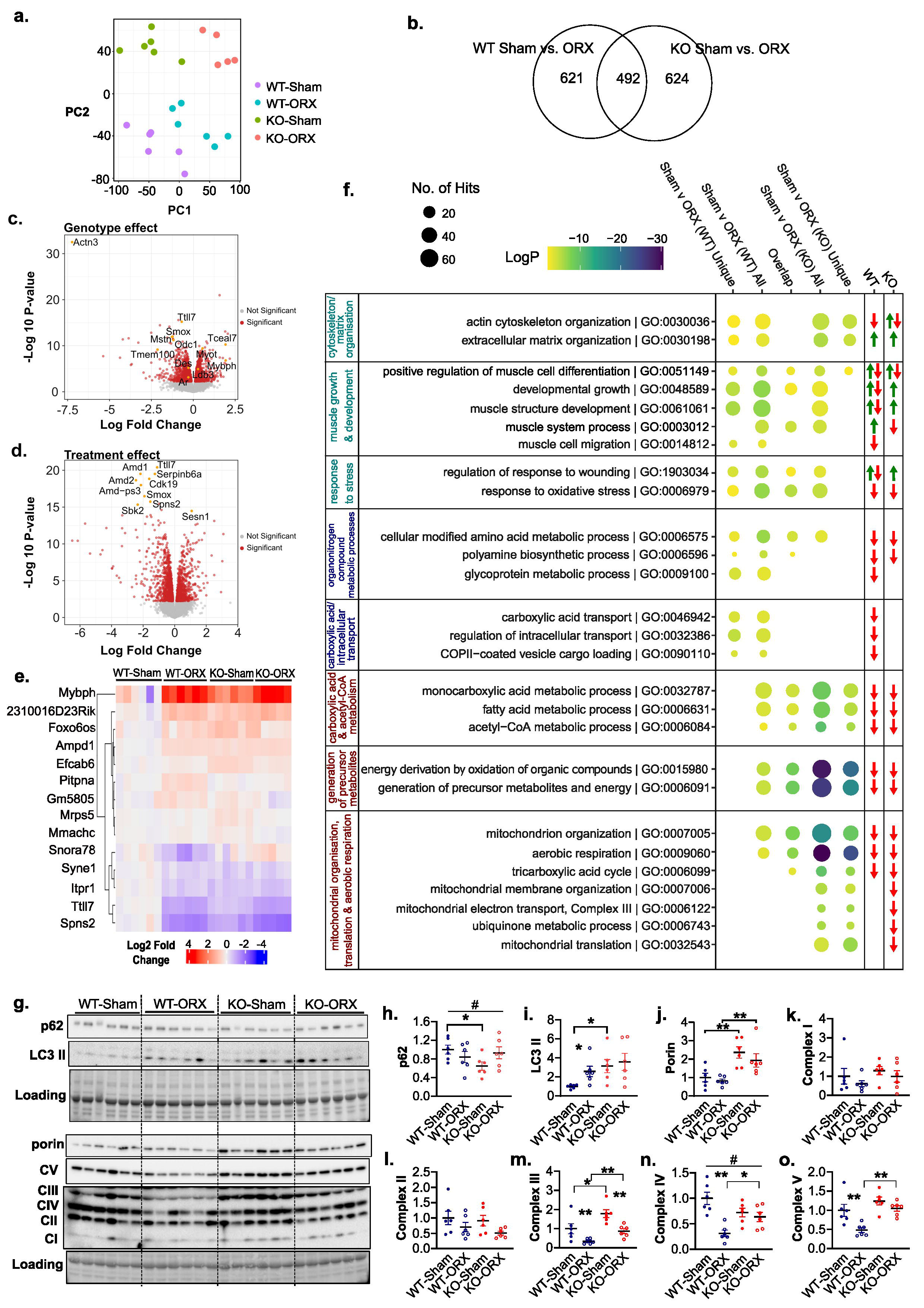
**Transcriptomic analysis highlights the differential metabolic response to androgen deprivation with α-actinin-3 deficiency. (**a) Principal Component Analysis (PCA) dot plot shows clear separation and grouping of samples by treatment (sham/orchidectomy) on PC1 and *Actn3* genotype (WT/KO) on PC2. (b) Venn diagram (*q* < 0.01) shows a comparable number of differentially expressed genes for WT and KO in response to orchidectomy, along with 492 genes that overlap for their effect in both genotypes compared to orchidectomy. (c-d) Volcano plots highlight the differentially expressed genes (*q* ≤ 0.01, in red) due to the effect of (c) *Actn3* genotype and (d) orchidectomy. (e) Heatmap shows expression in log2 fold-change of the 14 statistically significant genes (*q* < 0.05) from the interaction test performed. Fold changes are calculated against the mean of WT-Sham expression samples. (f) Summarised gene set enrichment analysis groups GO terms by commonality and illustrates the difference in response to orchidectomy between WT (WT unique, in blue) and *Actn3* KO (KO unique, in red) along with those that are present in both genotypes (in teal). The size of the circles represents the number of hits within the GO term and statistical significance in Log(P) is represented by the colour of the circle. Green and red arrows denote increased and decreased expression, respectively, of genes enriched in each GO term in ORX samples relative to Sham. (g) Western blot analysis confirms differential *Actn3* genotype effects on the response to orchidectomy for (h) autophagy marker p62 and (i) LC3 lipidation, as well as changes for (j) porin and (k-o) mitochondrial complexes I-V. Data are represented as mean ± SEM (h-o); \**p* < 0.05, \*\**p* < 0.01 by Mann-Whitney U test; #*p* < 0.05 by two-way ANOVA (h, n).

Tests of genotype effect (Fig 3c) confirm the transcript-level knockout of *Actn3* expression in *Actn3* KO relative to WT. As expected, we observe reduced expression of *Ar*, *Smox, Odc1, Tmem100 and Mstn*^26^ and increased expression of *Ldb3, Myot*^34^ and *Tceal7* with α-actinin-3 deficiency. Of note, *Mybph,* an androgen responsive gene^5,35,36^, is also upregulated with α- actinin-3 deficiency at baseline. Similarly, volcano plot of the treatment effect (Fig 3d) verifies the downregulation of the polyamine biosynthesis genes *Amd1* and *Smox* in response to orchidectomy among the top 10 genes that are differentially regulated in skeletal muscle^37^. Testing for effects of interaction between *Actn3* genotype and treatment (*q* < 0.05) identified 14 genes (Fig 3e), with the pattern of expression for these genes in KO groups showing remarkable similarity to WT-ORX.

We performed gene set enrichment analysis testing to further characterise the overall response to orchidectomy with respect to *Actn3* genotype. For each contrast, the top 15 most significant (by *q-*value) gene ontology (GO) terms were used to generate GO plots to examine both common and unique response to orchidectomy relative to Sham in WT and KO muscles (Supplementary Fig 3); this is summarised and organised in groups based on commonality using QuickGO in Fig 3f^38^. Arrows denote directionality of gene expression changes in ORX samples relative to Sham for genes enriched in each GO term, within each genotype. When examining the GO terms that are enriched in response to α-actinin-3 deficiency and orchidectomy, we observe that there are several overlapping pathways related to cytoskeleton/matrix organisation, muscle growth and development, response to stress, polyamine biosynthesis and cellular modified amino acid metabolism. The WT response to orchidectomy shows further unique enrichment for these pathways and demonstrates additional enrichment for GO terms relating to carboxylic acid/intracellular transport, which is not observed in the KO response. Both WT and *Actn3* KO muscles also show common responses for many GO terms related to the metabolism of carboxylic acid and acetyl-CoA, generation of precursor metabolites and mitochondrial metabolic processes, but *Actn3* KO demonstrate a greater amount of enrichment, as well as unique enrichment of other GO terms associated with mitochondrial activity. Interestingly, the expression of genes enriched in GO terms related to energy generation and metabolic processes is typically reduced in WT and KO in response to orchidectomy.

To verify a subset of these results, we examined specific markers of autophagy and mitochondrial activity in both WT and KO muscles, since they are associated with muscle atrophy induced by orchidectomy^39^, and autophagy is activated in response to oxidative stress and deficiencies in essential amino acids and ATP^40^ (Fig 3g-o). Contrary to our previous reports^26^, *Actn3* KO-Sham muscles show significantly higher lipidated LC3 (LC3-II), a standard marker for autophagosomes, and reduced p62 expression relative to WT-Sham muscles, consistent with increased autophagic flux with α-actinin-3 deficiency at baseline. Our results also show a significant interaction between the effects of *Actn3* genotype and orchidectomy on p62 expression (F(1, 20) = 4.457, *P* = 0.0475); WT-ORX show reduced p62 relative to WT-Sham, consistent with upregulated autophagy in WT following orchidectomy, while p62 levels showed a trend for increase in KO-ORX relative to sham control (Fig. 3h). A similar trend is observed for the expression of LC3-II (F(1,20) = 0.9108, *P* = 0.3513), with WT-ORX showing significantly increased LC3-II expression relative to WT-Sham (Fig. 3i), but not between KO-Sham and KO-ORX.

Assessment of VDAC/porin expression as a marker of mitochondrial number show no detectable difference in expression in either WT-ORX or KO-ORX relative to Sham controls, although porin expression is consistently higher in KO compared to WT (Fig. 3j). There is however a significant interaction between the effect of *Actn3* genotype and orchidectomy on the expression of Complex IV (F(1, 20) = 10.74, *P* = 0.0038). Consistent with previous reports, Complex IV expression is reduced with orchidectomy in WT muscles^39^. A similar trend is also observed for Complex V (F(1, 20) = 2.310, *P* = 0.1442) (Fig. 3k-o). Despite reduced transcript expression of genes related to aerobic respiration, protein expression of porin, Complex I, II, IV and V are not different between KO-ORX and KO-Sham. In sum, these results indicate that, in response to androgen deprivation, the absence of α-actinin-3 in skeletal muscle maintained the protein expression of mitochondrial complexes, while other pathways and processes that are typically activated in response to stress, such as autophagy, are suppressed.

### α-Actinin-3 deficiency reduces muscle hypertrophy in response to dihydrotestosterone (DHT)

We further examined the effect of *Actn3* genotype on the response to androgens in preventing muscle wasting and inducing muscle hypertrophy. Silastic tubing that are either empty or filled with crystalline dihydrotestosterone (DHT) were subcutaneously implanted in young, (aged 4-5 weeks, pre-pubertal) intact (surgical sham) and orchidectomised male mice, as well as female WT and *Actn3* KO mice for 6 weeks. LCMS analysis of sera analysis show similar increases of DHT, 5α-androstane-3α,17β-diol (3-α-diol) and 5α-androstane-3β,17β-diol (3-β-diol) (the two diols are the primary metabolites of DHT) in all WT and *Actn3* KO mice implanted with DHT relative to respective controls (Fig 4a, Supplementary Fig 4).

**Fig 4.**
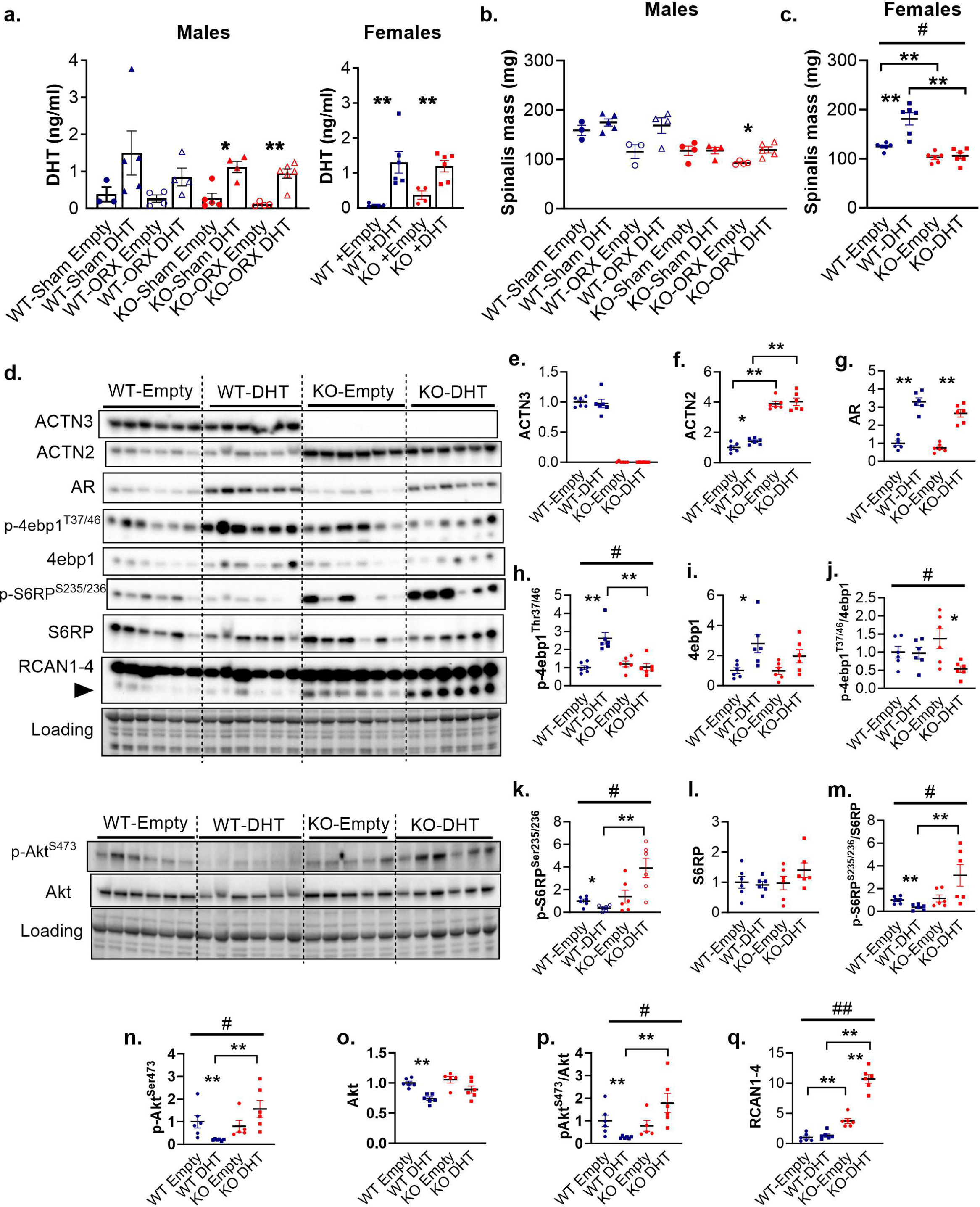
α-Actinin-3 deficiency decreases the DHT-induced muscle hypertrophic response. A) Liquid chromatography-mass spectrometry confirms increases in serum DHT in male (Sham/ORX) and female WT and *Actn3* KO mice that were implanted with silastic tubing containing 10 mg of crystalline DHT for 6 weeks, relative to mice that were implanted with empty tubing. b) DHT treatment prevented atrophy of spinalis muscles in orchidectomised WT and KO mice c) Female WT-DHT but not KO-DHT mice show increases in spinalis mass relative to empty controls. D) Western blot analysis quantify the effect of DHT in female WT and KO spinalis muscles on the expression of e) ACTN3, f) ACTN2, g) AR, h) p-4ebp1^Thr37/46^, i) total 4ebp1, j) ratio of p-4ebp1^Thr37/46^/4ebp1, k) p-S6RP^Ser235/236^, l) total S6RP, m) ratio of p- S6RP^Ser235/236^/S6RP, n) p-Akt^Ser473^, o) total Akt, p) ratio of p-Akt^Ser473^/Akt and q) RCAN1-4. Data are represented as mean ± SEM; \**p* < 0.05, \*\**p* < 0.01 by Mann-Whitney U test; # *p* < 0.05, ## *p* < 0.0001 by two-way ANOVA (c, h, j, k, m, n, p, q).

DHT treatment prevented the loss in overall body weight and muscle mass with orchidectomy in both WT-ORX and KO-ORX DHT mice compared to respective ORX mice implanted with empty controls and maintained seminal vesicle mass (Supplementary Table 4). The spinalis muscle, which is located adjacent to the site of implantation, show the greatest hypertrophic response to DHT compared to other muscles. There is a trend for lower % increase in spinalis mass in DHT treated *Actn3* KO-ORX than WT-ORX (F(1, 12) = 1.656, *P* = 0.2225; WT-ORX

DHT: +45.7%, *Actn3* KO-ORX DHT: +20.4%) (Fig 4b). Similar results are observed in female mice and two-way ANOVA confirms a significant *Actn3* genotype effect on DHT response for overall body weight (F(1, 20) = 4.928, *P* =0.0382), spinalis mass (F(1, 20) = 12.67, *P* = 0.0020) and heart mass (F(1, 19) = 9.791, *P* = 0.0055) (Supplementary Table 5). Relative to respective empty controls, female WT-DHT show greater increases than female KO-DHT for body mass (WT-DHT +20.36%, *P* =0.0022; KO-DHT +5.0%, *P* =0.4848), spinalis mass (WT-DHT: +45.2%, *P* =0.0022; *Actn3* KO-DHT: +3.17%, *P* = 0.6991) and heart mass (WT-DHT: +25.8%,*P* =0.0022, KO-DHT: +0.9%, *P* =0.9307) (Fig 4c, Supplementary Table 5). These results suggest that α-actinin-3 deficiency significantly reduces the overall growth response to DHT in female mice, with a similar trend observed in castrated male mice.

### α-Actinin-3 deficiency alters downstream Akt/mTOR and enhances calcineurin signalling in response to DHT

The expression of AR, α-actinins and downstream muscle protein synthesis signalling is further assessed in female spinalis muscles from empty- and DHT-treated WT and *Actn3* KO mice (Fig 4d-q). Following 6 weeks of DHT treatment, both WT-DHT and KO-DHT muscles show significant increases in AR expression compared to empty controls (Fig 4g). α-Actinin- 3 expression is unchanged with DHT in WT and KO (Fig 4e) but WT-DHT show a small significant increase in α-actinin-2 expression (Fig 4f). Evaluation of 4ebp1 and S6RP activation (as markers of protein synthesis) show significant interactions between the effect of *Actn3* genotype and DHT treatment on the expression of p-4ebp1^Thr37/46^ (F(1, 20) = 17.69, *P* = 0.0004) and p-S6RP^Ser235/236^ (F(1, 20) = 9.115, *P* = 0.0068), as well as ratio of p-4ebp1^Thr37/46^ /4ebp1 (F(1, 20) = 4.860, *P* = 0.0394) and p-S6RP^Ser235/236^/S6RP (F(1, 20) = 6.823, *P* = 0.0167).

There is also significant interaction between *Actn3* genotype and DHT effects for expression of p-Akt^Ser473^ (F(1, 19) = 8.645, *P* = 0.0084) and p-Akt ^Ser473^/Akt (F(1, 19) = 9.724, *P* = 0.0057).

Pair-wise comparisons with WT-empty show that WT-DHT muscles have significantly increased expression of p-4ebp1^Thr37/46^ and 4ebp1 (but not p-4ebp1^Thr37/46^/4ebp1) (Fig 4h-i) but show decreases in p-S6RP, p-S6RP^Ser235/236^/S6RP, p-Akt^Ser473^, Akt and p-Akt^Ser473^/Akt (Fig 4k- p). In contrast, KO-DHT show significant decreases in p-4ebp1^Thr37/46^/4ebp1, and a trend for increased p-S6RP^Ser235/236^ and p-S6RP^Ser235/236^/S6RP relative to KO-Empty.

We further examined changes in RCAN1-4 expression as a proxy for calcineurin activity. There is a highly significant differential *Actn3* genotype effect on RCAN1-4 in response to DHT (F(1, 20) = 51.97, *P* < 0.0001). RCAN1-4 expression is consistently higher in KO muscles compared to WT regardless of treatment, and while DHT treatment did not alter RCAN1-4 levels in WT muscles, RCAN1-4 is significantly increased in KO-DHT muscles relative to KO-Empty (Fig 4q).

### α-Actinin-3 deficiency significantly alters the transcriptional response to DHT

We further assessed global changes in the skeletal muscle transcriptome in spinalis muscles of female WT and *Actn3* KO mice treated with DHT. Principle component analysis demonstrates clear separation of samples based on DHT treatment and *Actn3* genotype (Fig 5a). DHT treatment results in significant differential expression (*q* ≤ 0.01) of 2140 genes in WT and 825 genes in *Actn3* KO muscles, consistent with a reduced response to DHT with α-actinin-3 deficiency; of these, 528 genes are common to both WT and *Actn3* KO (Fig 5b). Volcano plots of *Actn3* genotype (Fig 5c) and effects of DHT (Fig 5d) highlights many of the expected differentially expressed genes that are associated with these treatments^36^.

**Fig 5:**
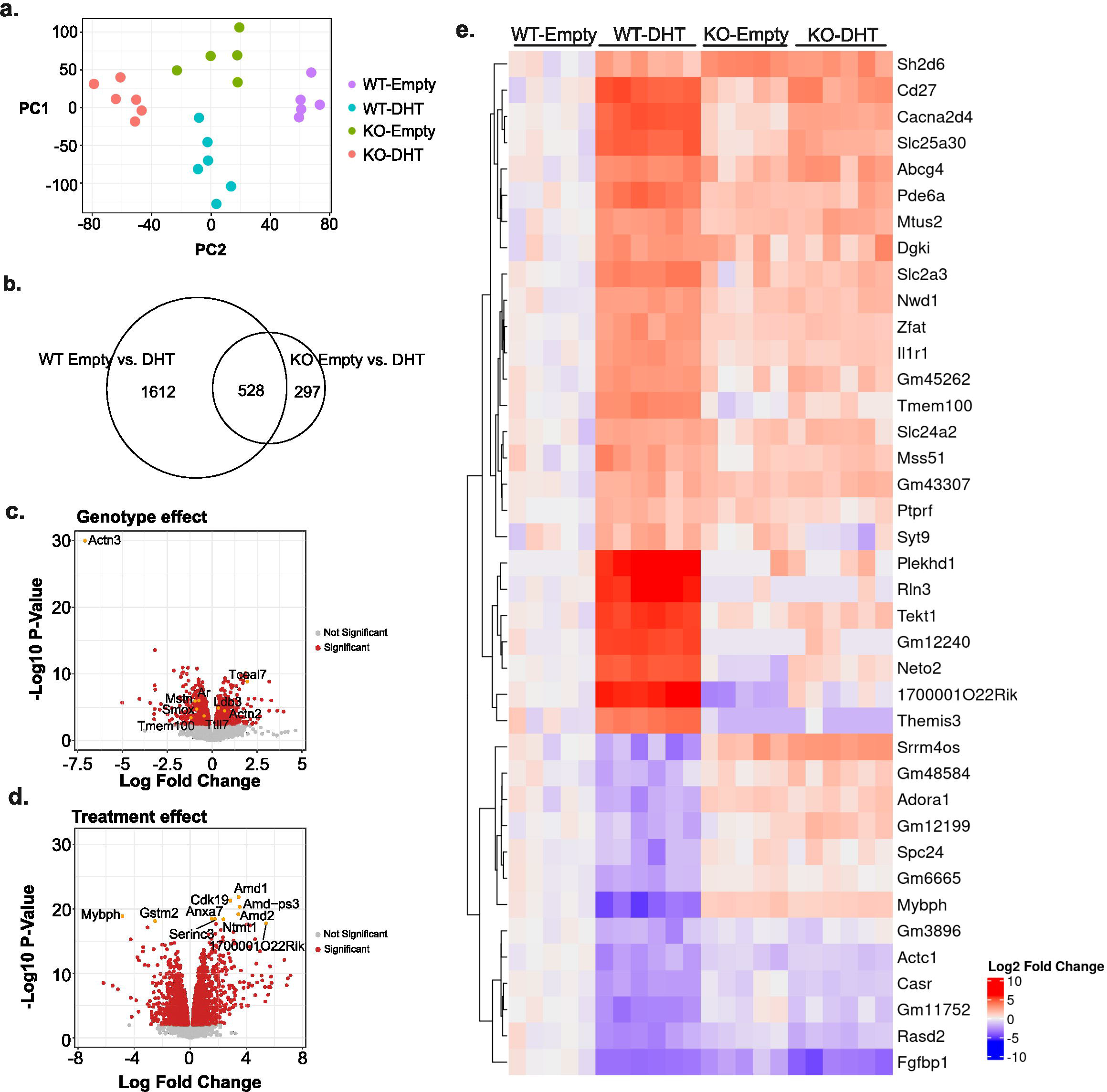
α-Actinin-3 deficient muscles show fewer differentially expressed genes in response to DHT. A) PCA plot shows clear separation and grouping of samples by *Actn3* genotype (WT/KO) and treatment (empty/DHT) in two dimensions. B) Venn diagrams (*q* < 0.01) illustrate greater differential gene expression for WT (1612 unique genes) in response to DHT compared to KO (297 unique genes), along with 528 genes that have an overlap of effect in both genotypes. (c-d) Volcano plots highlight the differentially expressed genes (*q* ≤ 0.01, in red) due to the effects of c) *Actn3* genotype and d) DHT. E) Heatmap shows the log2 fold change of genes that are statistically significant from the interaction between *Actn3* genotype and DHT treatment (*q* ≤ 0.01, log2 fold change ≥ 2). Fold changes are calculated against the mean of WT-Empty expression samples.

Testing for effects of interaction between *Actn3* genotype and DHT treatment (*q* ≤ 0.05) identified 624 genes; heatmap of gene expression changes of the top genes (*q* ≤ 0.01, log2 fold- change ≥ 2) shows minimal response to DHT in *Actn3* KO muscles (Fig 5e). To determine how WT and *Actn3* KO mice differentially respond to DHT, a gene set enrichment analysis is performed to examine both commonly activated and uniquely activated pathways based on *Actn3* genotype. The top 15 most significant GO terms for each contrast are recorded (Supplementary Fig 5); Fig. 6a shows a representative list organised into groups based on QuickGO^38^. Arrows denote directionality of gene expression changes in DHT samples relative to Empty for genes enriched in each GO term, within each genotype. In response to DHT, both WT and KO show commonality as well as unique gene sets associated with the regulation of cell junction and projection, oxidative stress response, transmembrane transport and muscle contraction and development. There is also an overlap between genotypes for processes relating to growth factor and autophagy response, amino acid and phospholipid metabolism, ribonucleotide synthesis and aerobic respiration. However, only WT show further enrichment for processes related to amino acid metabolism and modification, RAS/MAPK signalling and actin cytoskeleton organisation, while KO show further unique enrichment of GO terms and increased expression of genes for ribonucleotide synthesis and mitochondrial organisation and metabolism.

**Fig 6:**
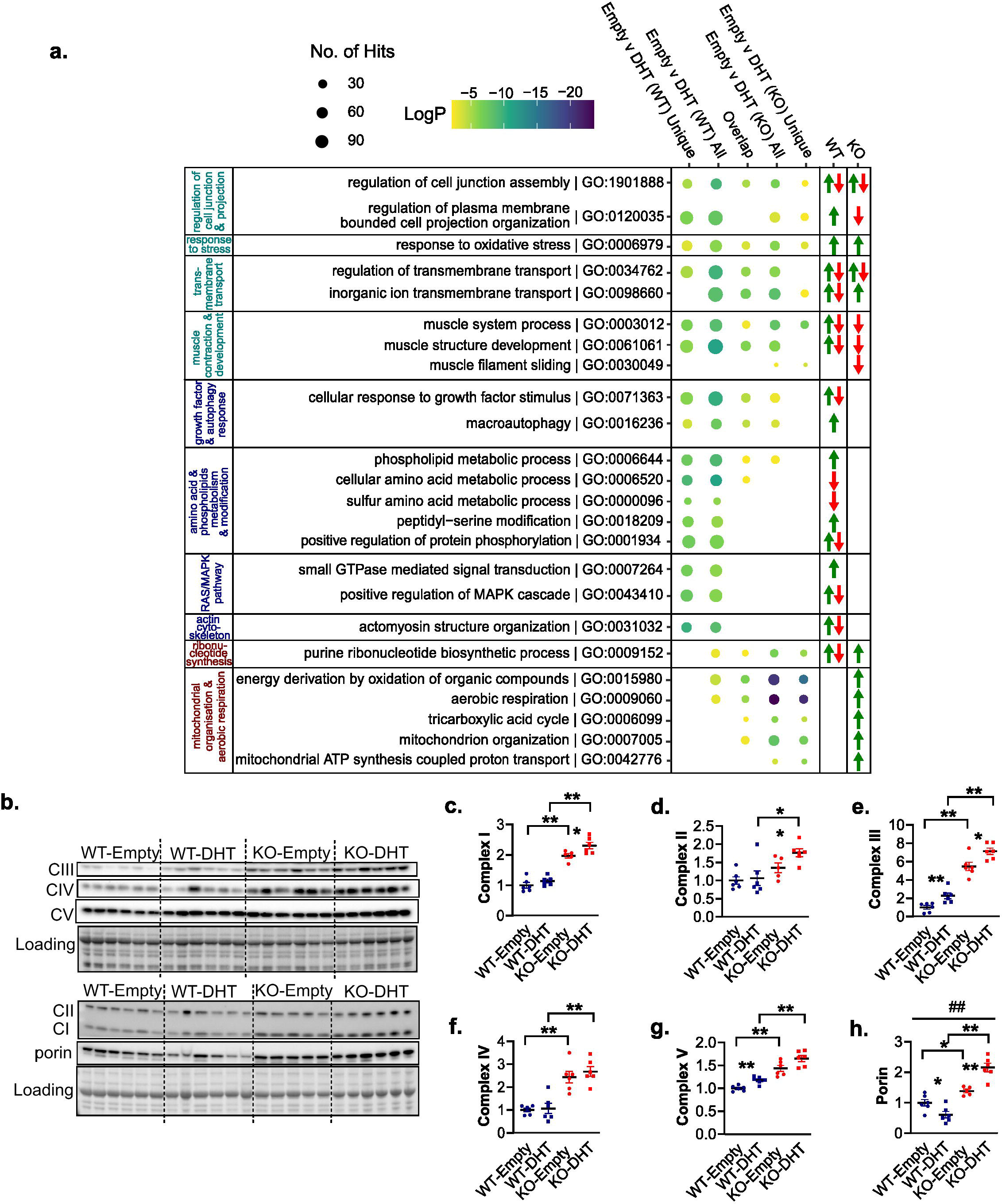
α-Actinin-3 deficiency alters the muscle transcriptomic response to DHT with preferential activation of mitochondrial metabolic processes. A) Gene set enrichment analysis was performed to compare the difference in DHT response between WT (WT unique) and *Actn3* KO (KO unique) along with those representing commonality. A representative list of the top GO terms for each contrast is shown. Groups in teal are common in WT and KO, groups in blue denote GO terms with unique enrichment in WT, while groups in red show further unique enrichment in KO. Green and red arrows denote increased and decreased expression, respectively, of genes enriched in each GO term in DHT samples relative to Empty. Western blot analysis confirms differential *Actn3* genotype effects on the response to DHT for mitochondrial complexes I-V (c-g) and (h) porin. Data are represented as mean ± SEM; \**p* < 0.05, \*\**p* < 0.01 by Mann-Whitney U test (c-h); ## *p* < 0.0001 by two-way ANOVA (h).

To confirm the differential activation of mitochondrial metabolism in response to DHT with α- actinin-3 deficiency, we further assessed the protein expression of mitochondrial complexes and VDAC/porin in WT and KO muscles (Fig 6c-i). KO muscles show higher expression of Complexes I-V and porin compared to WT, regardless of treatment. In contrast to the effects of orchidectomy, DHT treatment increased expression of Complex III and V in WT muscles relative to empty controls, while KO-DHT muscles showed significant increases for

Complexes I, II and III relative to KO-Empty. Testing for interaction effects between *Actn3* genotype and DHT treatment for expression of Complxes I-V did not reach statistical significance, but there was a significant interaction effect on the expression of porin (F(1,19) = 26.93, *P* <0.0001), with WT-DHT muscles showing reductions in porin while KO-DHT show increased expression relative to controls. Together, these results are consistent with an increase in mitochondrial number and activity with α-actinin-3 deficiency in response to DHT.

### α-Actinin-3-dependent mediators of androgen response in skeletal muscle

To determine the specific mediators of androgen response that are ACTN3-dependent, we examined the differentially expressed genes that show significant interaction between *Actn3* genotype and androgen deprivation, as well as *Actn3* genotype and DHT. A total of 8 genes are identified: *Mybph, 2310016D23Rik, Ampd1, Pitpna, Syne1, Itpr1, Ttll7* and *Spns2* (Fig 7a). The direction of change in expression of these genes relative to controls is inverse with androgen deprivation and DHT in both WT and KO samples, indicating these genes are androgen responsive. However, the magnitude of fold changes is smaller in KO relative to WT, suggesting that the androgen-dependent regulation of these genes also requires the presence of α-actinin-3, despite there being no reports of direct or indirect interactions of these gene products with the α-actinins.

**Fig 7.**
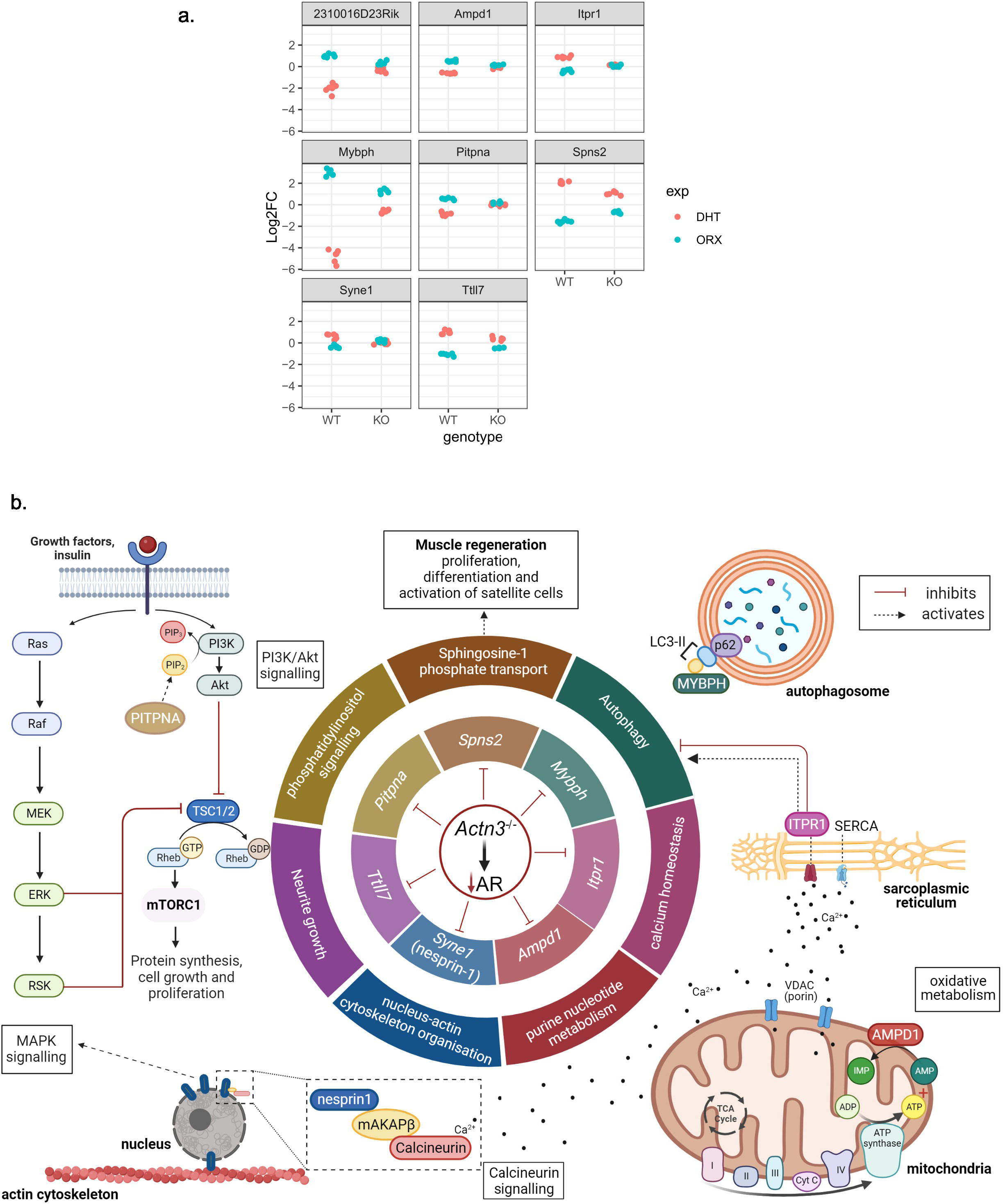
*α*-Actinin-3-dependent mediators of androgen response in skeletal muscle. a) Log2 fold-change (Log2FC) of differentially expressed genes (DEGs) that show significant interactions between orchidectomy and *Actn3* genotype, and also between DHT treatment and *Actn3* genotype. The DHT and ORX fold change response for these DEGs are inverse. For both experiments, gene expression changes are lower in *Actn3* KO samples. b) Schematic of proposed mechanistic changes with α-actinin-3 deficiency in response to androgen deprivation and DHT treatment. The absence of α-actinin-3 (which results in reduced muscle AR) inhibits/dampens the transcriptional response of *Mybph, Itpr1, Ampd1, Syne1, Ttll7, Pitpna and Spns2* to changes in circulating androgens. The putative functions of these genes relate to autophagy, calcium homeostasis, purine nucleotide metabolism, nucleus-actin cytoskeleton organisation, neurite growth, phosphatidylinositol signalling and sphingosine-1 phosphate transport, respectively. The altered response of these genes to androgen deprivation and DHT treatment with α-actinin-3 deficiency may contribute to many of the differential downstream signalling (eg. Calcineurin, MAPK, PI3K/Akt/mTORC1, autophagy and oxidative metabolism) that influence muscle mass. The gene *2310016D23Rik* is also differentially expressed but is excluded from the schematic since its function is unknown. (created with BioRender.com)

The putative functions of these genes vary greatly, ranging from autophagy (*Mybph*), calcium homeostasis (*Itpr1*), purine nucleotide metabolism (*Ampd1*), phosphatidylinositol signalling (*Pitpna*), neurite growth (*Ttll7*), sphinogosine-1 phosphate transport (*Spns2*) and nucleus-actin cytoskeleton organisation (*Syne1*). With the exception of *Ttll7* and *2310016D23Rik,* these genes have also been associated with muscle atrophy and disease (Supplementary Table 6). The functions of these genes correlate with many of the pathways identified in Fig 4f and 6a, suggesting that the dampened transcriptional response of these genes may have mediated or contributed to many of the differential signalling changes with α-actinin-3 deficiency in response to androgen deprivation and DHT treatment (Fig 7b).

## Discussion

In this study, we show for the first time that homozygous inheritance of a common null polymorphism in the *ACTN3* gene results in significantly reduced muscle androgen receptor (AR) protein expression in humans and mice, in both males and females. This finding is remarkable since muscle AR has been shown to be a key regulator of muscle force production, fatigue resistance, sarcomeric organisation and fibre type distribution in various male ARKO models^5,25,36,41–45^ and androgen sensitivity in skeletal muscle is strongly correlated with AR expression^46^. Given that α-actinin-3 deficiency is common in the general population, our results suggest that healthy adult human muscle function is sustainable with ∼70% reduction in functional AR in muscle, albeit at some cost to sprint and power performance.

The reduction of AR in the testis of *Actn3* KO mice also had no effect on baseline levels of serum testosterone, luteinising hormone and seminal vesicle mass indicating no specific effects on testosterone or sperm production^47^. Fertility in *Actn3* KO mice appears normal; however, testis size is marginally but significantly increased. This may be an adaptive response to maintain serum testosterone. In support of our findings in mice, a recent genome wide association study of 425,097 UK Biobank study participants (aged 40-69 years) did not identify *ACTN3* rs1815739 as a genetic determinant of testosterone levels and related sex hormone traits in men and women^48^, contrasting the previous report of an association between increased serum testosterone with the *ACTN3* 577R allele in elite Russian athletes^20^. Overall, these data suggest that the phenotypic effects of α-actinin-3 deficiency in skeletal muscle are a direct consequence of reductions in AR and not of changes in circulating testosterone in the general population.

Our findings indicate that the effect of α-actinin-3 deficiency is specific to AR and does not impact ER and TR signalling, despite α-actinin-2 (which is upregulated with α-actinin-3 deficiency) being a co-activator for these nuclear receptors^9^. We also show that α-actinin-3 is directly and positively associated with AR expression in mammalian skeletal muscle, indicating that α-actinin-3 plays a specialised role in the regulation of AR signalling that cannot be compensated by increased expression of α-actinin-2^11^, even though they share 80% amino acid sequence identity^49^. These results complement earlier findings of reduced *Actn3* (but not *Actn2*) expression in testicular feminised mice that lack functional androgen receptors in the testis^21^, and in Sertoli cell-specific ARKO mice^50^, confirming the presence of a specific and direct relationship between α-actinin-3 with AR signalling.

Consistent with our hypothesis, decreased baseline AR with α-actinin-3 deficiency alters the muscle adaptive response to changes in circulating androgens. Both female and castrated male *Actn3* KO mice showed lower levels of muscle hypertrophy following DHT treatment. α- Actinin-3 deficiency also enhanced the atrophic response to androgen deprivation in the gastrocnemius (a highly androgen-sensitive muscle), although the overall whole-body response to orchidectomy was not different from controls. The differential response in muscle adaptation to changes in androgen levels with α-actinin-3 deficiency is accompanied by differences in the activation of downstream mTORC1 and calcineurin-NFAT signalling. Both WT and *Actn3* KO muscles show a similar trend for reduced phosphorylation of mTORC1 substrates 4ebp1 and S6RP with androgen deprivation, but there was a genotype difference in the phosphorylation of these substrates in response to DHT. RCAN1-4 expression (as a marker of calcineurin signalling) is unchanged in WT muscles in response to DHT treatment and androgen deprivation, however in *Actn3* KO muscle, RCAN1-4 is significantly increased with DHT treatment and markedly reduced in response to androgen deprivation.

It is unclear the extent to which differences in mTORC1 and calcineurin activation account for the increased atrophic response to androgen deprivation and reduced hypertrophic response to DHT in *Actn3* KO mice. The involvement of these pathways in muscle atrophy/hypertrophy in response to androgen deprivation and testosterone treatment, respectively, is widely disputed, with recent studies suggesting that mTORC1 has a limited role in testosterone-induced muscle growth^4,23,51–55^. The lack of change in RCAN1-4 expression in WT-ORX and WT-DHT muscles in our study is also consistent with others that do not support a role for calcineurin in muscle growth^56^. However, our finding of inverse responses in RCAN1-4 expression with androgen deprivation and DHT treatment that is specific to *Actn3* KO muscles strongly suggests that calcineurin signalling is involved in muscle adaptation to changes in testosterone in the absence of α-actinin-3 and reduced AR. We have previously reported increases in calcineurin activity in muscles from *Actn3* KO mice and in *ACTN3* 577XX humans at baseline, and upregulation of this pathway in *Actn3* KO mice enhances the response to exercise training, modifies the muscle adaptive response to denervation and immobilisation and slows the progression of muscular dystrophy in *mdx* mice^24,26,28,57^. These results further confirm calcineurin signalling as a critical pathway that is preferentially activated in the absence of α- actinin-3 during muscle remodelling.

Hypothesis-free transcriptomic analyses provided additional insights into the molecular pathways that are differentially altered with α-actinin-3 deficiency in response to changes in androgens. Heatmaps of the differentially expressed genes with *Actn3* genotype and orchidectomy or DHT treatment clearly illustrate a lack of change in gene expression in *Actn3* KO compared to WT, confirming that androgen sensitivity is reduced in α-actinin-3 deficient muscles. Comparison of the expression profiles of the significant interaction genes in male KO-Sham and WT-ORX muscles also show remarkable similarity, suggesting that α-actinin-3 deficient muscles in some respects are already in a “pre-castrated” state. Indeed, muscle AR expression levels of KO-Sham are similar to that of WT-ORX, and this could underpin the lower body mass and lean mass seen with α-actinin-3 deficiency at baseline.

Results from our gene set enrichment analyses show that androgen deprivation in male mice and DHT treatment in prepubertal female mice commonly alter expression of genes involved in oxidative metabolism, amino acid catabolism, oxidative stress, muscle growth and development, regardless of genotype. Myofiber AR expression has been shown to directly influence expression of these genes^36,45,58^, hence the coactivation/repression of these processes in response to changes in circulating androgens is consistent with published reports. Orchidectomy has been shown to reduce AR expression and downstream mTORC1 protein synthesis signalling as well as activate pathways that regulate autophagy and mitochondrial activity^54,55,59^. In contrast, DHT increases cell proliferation, ATP production, amino acid transport and protein synthesis through the epidermal growth factor receptor and MAPK activation^60,61^.

Our finding that WT and KO show differential enrichment of these pathways in response to orchidectomy and DHT suggest that transcription of many of these genes is also α-actinin-3 dependent, in addition to being sensitive to changes in androgens. For example, GO terms related to the MAPK cascade, amino acid metabolism and modification are only enriched in WT-DHT muscles, suggesting that α-actinin-3 expression is necessary for activating transcription of these genes, consistent with the suppression of muscle hypertrophy in *Actn3* KO mice in response to DHT treatment. Similarly, processes related to carboxylic acid/intracellular transport, glycoprotein metabolism are only altered in WT-ORX. This is further supported by the differentially expressed genes that show significant interaction with *Actn3* genotype (Fig. 7, Suppl Table 6) whose functions correspond to these processes. In sum, our data strongly suggest that α-actinin-3 is a key molecular partner in the coupling of these typical downstream signalling response to changes in androgens.

Among these, myosin binding protein H (*Mybph;* MyBP-H) is of particular interest. MyBP-H is primarily expressed in fast-twitch muscle fibres and has a purported role in autophagy processes in cardiomyocytes as it colocalises with LC3 at the autophagosome membrane^62,63^. *Mybph* expression is higher in gastrocnemius muscles of male global ARKO mice^23^, and is up-regulated in response to castration and down-regulated with testosterone/DHT treatment^5,35,36^. Interestingly, *Mybph* is also consistently among the top genes upregulated with α-actinin-3 deficiency at baseline in male (Fig 3c) and female mice (*q* = 0.09, data not shown). On this basis, persistent upregulated *Mybph* could explain why muscle mass is reduced with α-actinin- 3 deficiency. It also indicates that α-actinin-3 directly regulates *Mybph* transcription, and thus may play an indirect role in autophagic activation.

Conversely, GO terms related to mitochondrial organisation and activity are preferentially enriched with α-actinin-3 deficiency, both in response to orchidectomy and DHT treatment. Evaluation of OXPHOS complexes shows reduction of complexes III, IV and V in WT-ORX muscles, while expression levels of complex IV and V are largely maintained in KO-ORX. In contrast, DHT treatment significantly increased porin expression in KO muscles, but reduced porin levels in WT-DHT relative to controls. It is likely that mitochondrial remodelling could be a primary adaptive mechanism in skeletal muscles in the absence of α-actinin-3 and reduced AR, since *ACTN3* replacement (“rescue”) in *Actn3* KO muscles also specifically reduces oxidative metabolism, resulting in reduced force recovery after fatigue^17^. Taken together, our findings indicate that the absence of α-actinin-3 results in a failure to coactivate many of the typical pathways in response to changes in androgens, leading to a reliance on leveraging mitochondrial remodelling, and calcineurin signalling to restore muscle homeostasis. Our results also highlight the pivotal role of α-actinin-3 in various processes associated with the regulation of protein turnover and muscle mass.

Finally, it is important to note that our findings in the *Actn3* KO mouse are likely the aggregate effects of a global reduction in AR, and not merely a consequence of reduced muscle AR. Data from GTEX indicate that the *ACTN3* 577XX genotype in humans is associated with a loss of *ACTN3* transcript expression in other tissues (data not shown), suggesting that AR protein is likely reduced globally. We have shown that AR protein expression is reduced in skeletal muscles and testes of *Actn3* KO mice. This is an important distinction, since muscle-specific ARKO mice do not display differential changes in hindlimb muscle mass following androgen deprivation or DHT treatment^36,41^. Similarly, in our rAAV-mediated *ACTN3* gene “rescue” study^17^, we did not observe muscle hypertrophy, and this was likely due to the increase of AR being muscle-specific. Others have demonstrated that AR levels in neurons may in fact play a greater role in regulating muscle mass^64^. This raises the possibility that globally increasing α- actinin-3 expression (and thus AR) may increase muscle mass.

## Conclusion

In conclusion, we have demonstrated for the first time that α-actinin-3 deficiency is associated with reduced AR protein expression in skeletal muscles and testes, and influences androgen sensitivity in skeletal muscle. We have also shown that α-actinin-3 plays a distinct role in regulating the expression of genes in various pathways that influence muscle growth and metabolism. Our findings suggest that changes in mitochondrial activity are the primary adaptive response of α-actinin-3 deficient muscle to changes in androgen levels, consistent with the hypothesis that the *ACTN3* 577X allele was positively selected during recent human evolution due to enhanced muscle metabolic efficiency^11^.

Furthermore, our results suggest that *ACTN3* R577X genotype will contribute to the clinical variability in patients with partial or mild androgen insensitivity syndrome, and may also affect the health outcomes and treatment response of patients with prostate cancer who undergo androgen deprivation therapies (ADT)^65^. *Actn3* KO mice demonstrate increased muscle wasting following androgen deprivation, particularly in the weight-bearing gastrocnemius muscle, which is highly susceptible to atrophy in sarcopenic and cachexic conditions^35,66,67^, suggesting that α-actinin-3 deficient patients receiving ADT may be at increased risk of muscle wasting. In addition, reduced hypertrophic response to DHT in *Actn3* KO mice suggests that testosterone replacement therapy in isolation may be less effective for inducing fat free mass accretion and increases in muscle strength in α-actinin-3 deficient patients (eg. HIV-infected men)^68,69^ and in athletes who engage in androgen doping.

## Materials and Methods

### Animals and Ethics

All experiments were performed in male and female C57BL/6J mice. Mice were fed standard chow and water *ad-libitum* and were maintained in a 12:12 hour cycle of light and dark at ambient room temperature (∼22°C). All experiments were approved by the Animal Ethics Committee of the Murdoch Children’s Research Institute (A917).

### Human muscle biopsies

Protein was extracted from human muscle biopsies obtained from female and male participants for analysis of AR expression. Female participants (*n =* 33) were healthy, premenopausal aged 18-37; *n* = 23 were not resistance trained and *n =* 10 were resistance trained. Male participants (*n* = 11) were also a combination of healthy sedentary and resistance trained subjects and were aged 20-40. RNA was also obtained from vastus lateralis muscles of a separate cohort of moderately trained Caucasian men (aged 18-47; *N* = 24) and women (aged 21-45; *N* = 20)^70^ for evaluation of *AR* gene expression by qPCR.

### RT-PCR

cDNA was generated using 5 ng of total RNA using iScript Reverse Transcriptase supermix (cat#1708841) and C1000 Touch Thermal Cycler (Biorad) as per manufacturer guidelines. RT-qPCR was performed in triplicate with the LightCycler 480 instrument (Roche Diagnostics) and LightCycler 384-well plates with sealing foil (Roche Diagnostics). Reaction volume of 10 μl contains 2× SensiFAST SYBR No-ROX mix (Bioline), 0.4 μl of each 10 μM forward and reverse primers, and 1 μl of cDNA. Amplification of the single PCR product was confirmed using the melting point dissociation curve, and the *Cp* values were calculated using the LightCycler 480 software. For each gene, a standard curve was generated using serial dilution of plasmid DNA. Gene expression is then normalized to the geomean of two housekeepers *RPL27* and *RPL41*. Primers for RT-PCR reactions are as follows: *ACTN3* (forward: GACAGCTGCCAACAGGATCT and reverse: ATCCACTCCAGCAGCTCACT), *AR* (forward: CTTCGCCCCTGATCTGGTTT and reverse: CTCATTCGGACACACTGG-CT), *RPL27* (forward: GCAAGAAGAAGATCGCCAAG and reverse: TCCAAGGGGATAT- CCACAGA), *RPL41* (forward: AAGTGGAGGAAGAAGCGAATG and reverse: TGGACCTCTGCCTCATCTTT).

### ACTN3 over-expression studies

rAAV vectors containing human *ACTN3* (rAAV-CMV-*ACTN3*) were delivered by intramuscular injection into the anterior compartment of the hindlimb of anaesthetised *Actn3* KO mice, as previously described^17^. An injection containing 5x10^8^-5x10^10^ vector genomes of r-AAV-CMV-*ACTN3* in 30 μl of HBSS was delivered throughout the length of the tibialis anterior (TA) muscle. The contralateral limb was given an equivalent vector genomes of empty control vector. Mice were euthanised 6 weeks post injection and the TA muscles harvested for analysis.

### Dual X-ray Absorptiometry (DXA)

Changes in body composition, including lean body mass, fat mass, bone mineral density (BMD) and bone mineral content (BMC) were assessed using GE Lunar Piximus2 DXA scanner to assess mice 12 weeks post orchidectomy. All calculations were performed excluding the head and tail of the mice.

### Orchidectomy

Male mice aged between 8-10 weeks were given pre-surgical analgesia (buprenorphine 0.1mg/kg) and anaesthetised using isoflurane/oxygen. Incision was made via the abdomen to remove the fat pad, testes and vas deferens; the vas deferens was sutured and wound site was resealed with surgical clips for 10 days. Control animals received a sham orchiectomy, whereby the testes were surgically exposed but not cut and the wound site sealed with surgical clips. Animals were monitored and sacrificed 12- weeks post-surgery.

### Implantation with dihydrotestosterone (DHT)

Silastic tube implant prepacked with 10 mg of solid DHT or empty tubing^71^ were subcutaneously implanted between the scapulae in male and female mice aged 4-5 weeks. Mice were given pre-surgical analgesia (buprenorphine 0.1mg/kg) and fully anaesthetised using isoflurane/oxygen during implantation. Male mice were also orchiectomised at the same time to eliminate contributions from endogenous androgens. Mice were euthanised after 6 weeks of treatment.

### Steroid Profile Testing

The steroid profiles from sera including androgens, estrogens; testosterone (T), dihydrotestosterone (DHT), 5α-androstane-3α,17β-diol (3αDiol), 5α-androstane-3β,17β-diol (3βDiol), estradiol (E2) and estrone (E1) were assessed by liquid chromatography–tandem mass spectrometry (LC–MS/MS) assays (The ANZAC Research Institute)^72^.

### Immunoblotting

Snap frozen gastrocnemius or spinalis muscles were homogenised in 4% SDS lysis buffer and assessed for total protein concentration using Direct Detect Assay free cards (Millipore, Cat #DDAC00010-GR). Proteins were separated by SDS-PAGE using stain-free precast midi- criterion gels (Biorad), then transferred to polyvinylidene fluoride membranes (PVDF, Millipore). These were blocked with 5% BSA in 1× TBST, probed overnight at 4°C with primary antibodies against α-actinin-3, α-actinin-2 (ab68204, ab68167, abcam), Akt1 (#C73H10, CST), p-Akt (Ser463) (#4060, CST), mTOR (#2983, CST), p-mTOR (#5536, CST), p-4ebp1 (#2855, CST), 4ebp1 (#9452, CST), S6RP (#2217, CST), p-S6RP (#4856, CST), SQSTM1/p62 (#5114, CST), LC3B (#3868, CST), p-AR (ab45089, abcam). Blots were developed with Clarity Western ECL substrates (Biorad) using Bio-rad chemidoc. Densitometry was performed using Bio-Rad ImageLab software. Results were normalized to total protein and presented relative to WT control.

### Fibre morphometry analysis

Fibre typing was performed as previously described^73^. Sections were imaged on the V-Slide Scanner (MetaSystems) and analysed using Metamorph software (Molecular Devices).

### RNA-seq analysis

Total RNA was extracted from ∼ 50 mg of mouse gastrocnemius (androgen deprivation experiment) or spinalis (DHT treatment experiment) by phenol chloroform extraction (1 mL TRIsure solution; Bioline), purified using the RNeasy Mini Kit (Qiagen) and RNA integrity was determined using TapeStation (Agilent Technologies 2200). Libraries were prepared using the Illumina TruSeq Stranded Total RNA GOLD and sequenced on an Illumina NovaSeq 6000. Each sample was sequenced at 2x150bp at approximately 40M reads. Sample fastq files were processed using the RNASik pipeline^74^. Sample reads were aligned using STAR^75^, marked duplicates using PICARD^76^ and expression quantified reads with featureCounts^77^. Differential gene expression analysis was performed in Degust (version 4.11) and R-4.3.2 *limma*^78^. Lowly expressed genes were removed with a cutoff of 0.5 CPM in at least two samples. *P*-values were adjusted using the Benjamini-Hochberg method. As the experiment was structured as a pair of two factor experiments, genotype and treatment effects were probed separately and then compared by overlapping sets of differentially expressed genes. Interaction effects were modelled as a difference of differences. Gene set enrichment analysis was performed with Metascape^79^ and grouped using QuickGO^38^. To visualise data, figures were generated using R- 4.3.2 ggplot2 and ComplexHeatmap^80^. Directionality of GO terms were established by performing a second round of GO term enrichment of differentially expressed genes that were sorted by foldchange into “Up” and “Down” sets and observing from which set the GO term is enriched.

### Statistics

Analyses were performed in Graphpad Prism (V7, Graphpad Software Inc.) and StataSE (StataCorp). As group sizes consisted of <12, two-sided, unpaired t-tests using nonparametric statistics, Mann-Whitney U-tests were applied using an alpha of 0.05 for all analyses. Two-way ANOVA was applied to determine effects of genotype on interventions. Data were presented as individual points with the calculated group mean (line), or as bar graphs, with ±SEM error bars for each group.

## Acknowledgements

This work was supported by the Australian National Health and Medical Research Council (NHMRC) project grant (APP1130215) awarded to J.T.S. and K.N.N., and by a research grant from the World Anti-Doping Agency (16E11FP) awarded to Y.P.. K.N.R. was supported by a NHMRC Dora Lush Postgraduate Scholarship (GNT1114935). Se.L. was supported by a Future Fellowship from the Australian Research Council (FT210100278). D.J.H. is supported by a NHMRC L3 Investigator Grant (#1197361). The Novo Nordisk Foundation Center for Stem Cell Medicine, reNEW, is supported by a Novo Nordisk Foundation grant number NNF21CC0073729.

## Author Contributions

K.N.R. performed the baseline, castration and DHT experiments in mice. J.T.S. and K.N.R. performed the mouse western blots and generated the figures. M.S., F.J.R., M.R. and H.D.W. performed the transcriptomics analyses. L.R.M. assessed AR levels with ACTN3 over- expression and performed the fibre morphometry analysis. G.L., A.K.T., Y.P., Sh.L., S.E.A., Se.L., N.E. supplied human samples and performed the western blots in human skeletal muscles. W.C. and T.M. performed the DEXA analysis. C.T., P.H. assisted with tissue harvesting. D.H. supplied the DHT and performed the steroid profile testing. J.T.S. and K.N.N. designed the study. J.T.S. interpreted the data and wrote the paper. All authors provided feedback on the final draft of the paper.

## Conflict of Interest

F.J.R. receives institutional support as a coinvestigator and subcontracted by the Peter MacCallum Cancer Centre for an investigator-initiated trial which receives funding support from Sanofi/Regeneron Pharmaceuticals. The World Anti-Doping Agency did not have a role in the design of the study or in the collection, analysis and interpretation of data or the writing of the manuscript.

## Supplementary Figure Legends

**Supplementary** Fig 1. a) qPCR analysis of *AR* in vastus lateralis muscles from a small cohort of moderately trained Caucasian men (aged 18-47; *N* = 24) and women (aged 21-45; *N* = 20) showed a trend for reduced *AR* expression in 577XX individuals compared to RR+RX. b) p- AR^Ser213^ is upregulated in *Actn3* KO muscles compared to WT. c) ER-β protein expression in female gastrocnemius muscle are similar between WT and *Actn3* KO. Expression levels are normalised to actin. d) Immunostaining of ER-α shows similar staining intensities and localisation in female WT and *Actn3* KO muscles, scale bar = 50 μM. e) eQTL scan in the GTEx testis dataset show that the R577X variant is significantly associated with *ACTN3* transcript expression *P* = 8.93 × 10^−51^). f) Circulating levels of luteinising hormone (LH) were determined by radioimmunoassay assay and are not significantly different between male WT (*n* = 11) and *Actn3* KO (*n* = 10) mice. Data are represented as mean ± SEM. \*\**p* < 0.01 by Mann-Whitney U test.

**Supplementary** Fig 2. Fibre proportions in the gastrocnemius muscle of SHAM and ORX WT and *Actn3* KO mice. Fast fibre type proportions are not different in WT or *Actn3* KO muscles in response to castration, however slow type 1 proportions were marginally increased in WT. *N*=6-13 animals were analysed in each group. Data are represented as mean ± SEM. \*\*\**p* < 0.001 by Mann-Whitney U test.

**Supplementary** Fig 3. The top 15 gene ontology (GO) terms (by *q-*value) from each of the 5 contrasts demonstrate both common and unique response to orchidectomy relative to Sham in WT and KO muscle tissue samples.

**Supplementary** Fig 4. Liquid chromatography-mass spectrometry analysis of 3-α-diol and 3-β-diol levels (the primary DHT metabolites) in male (a, b) and female (c, d) mice. Data are represented as mean ± SEM; \**p* < 0.05, \*\**p* < 0.01 by Mann-Whitney U test.

**Supplementary** Fig 5. Dot plot of the top 15 most significant GO terms for each assessed contrast to investigate both common and unique responses to DHT relative to Empty in WT and KO muscles.

