## Supplementary figures and images for "*ACTN3* genotype influences androgen response in skeletal muscle"

### Supplementary Fig 1

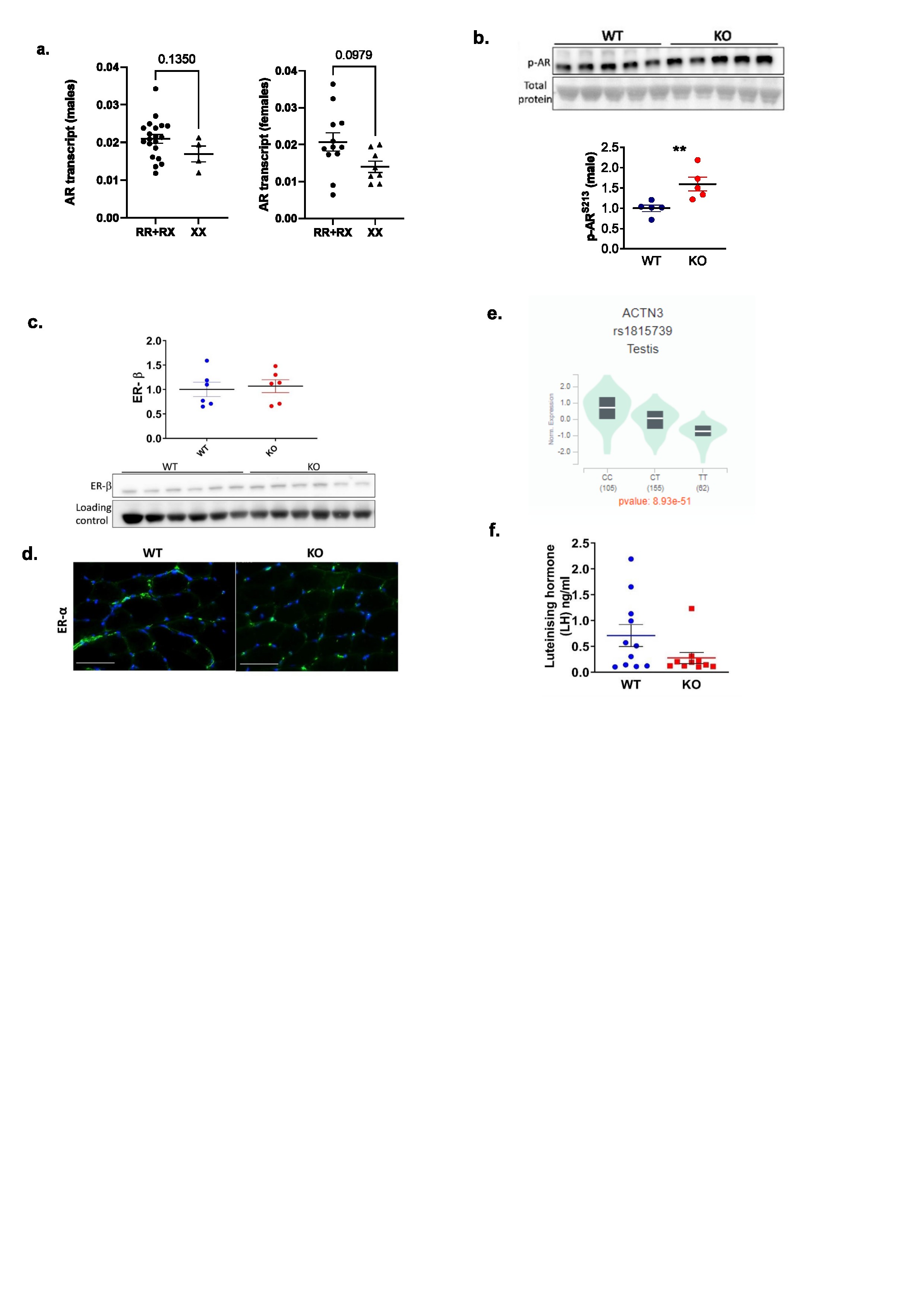

### Supplementary Fig 2

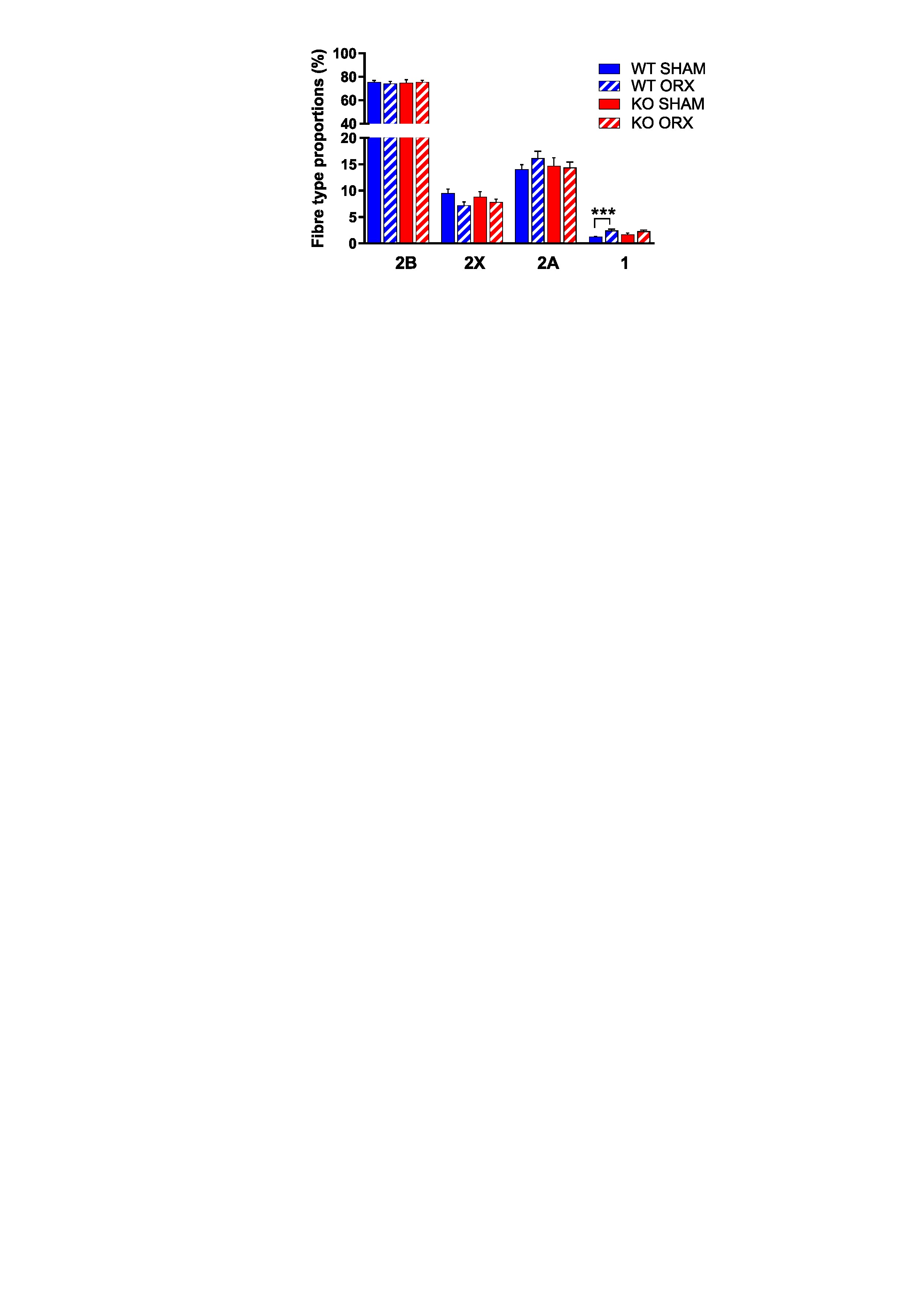

### Supplementary Fig 3

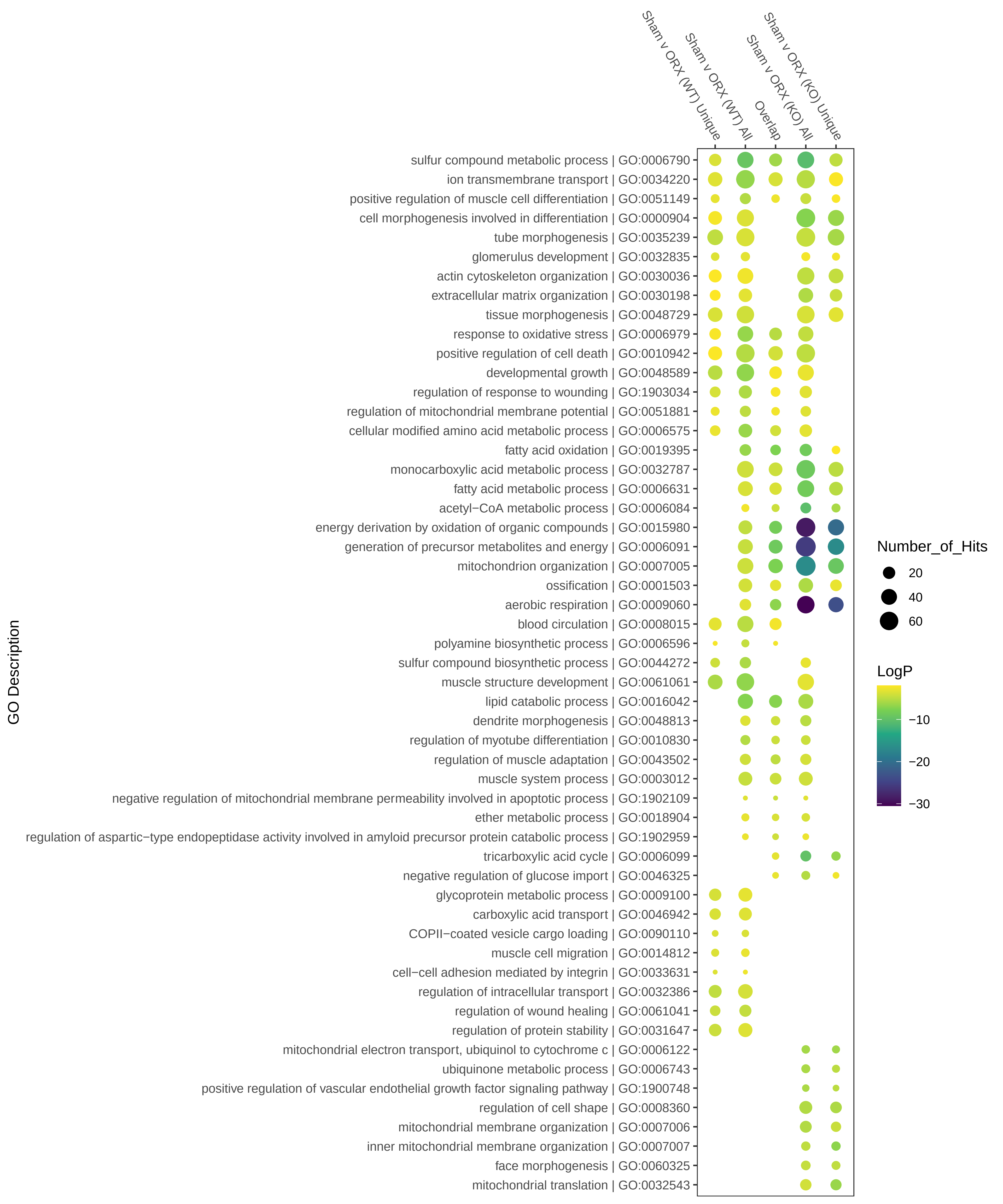

### Supplementary Fig 4

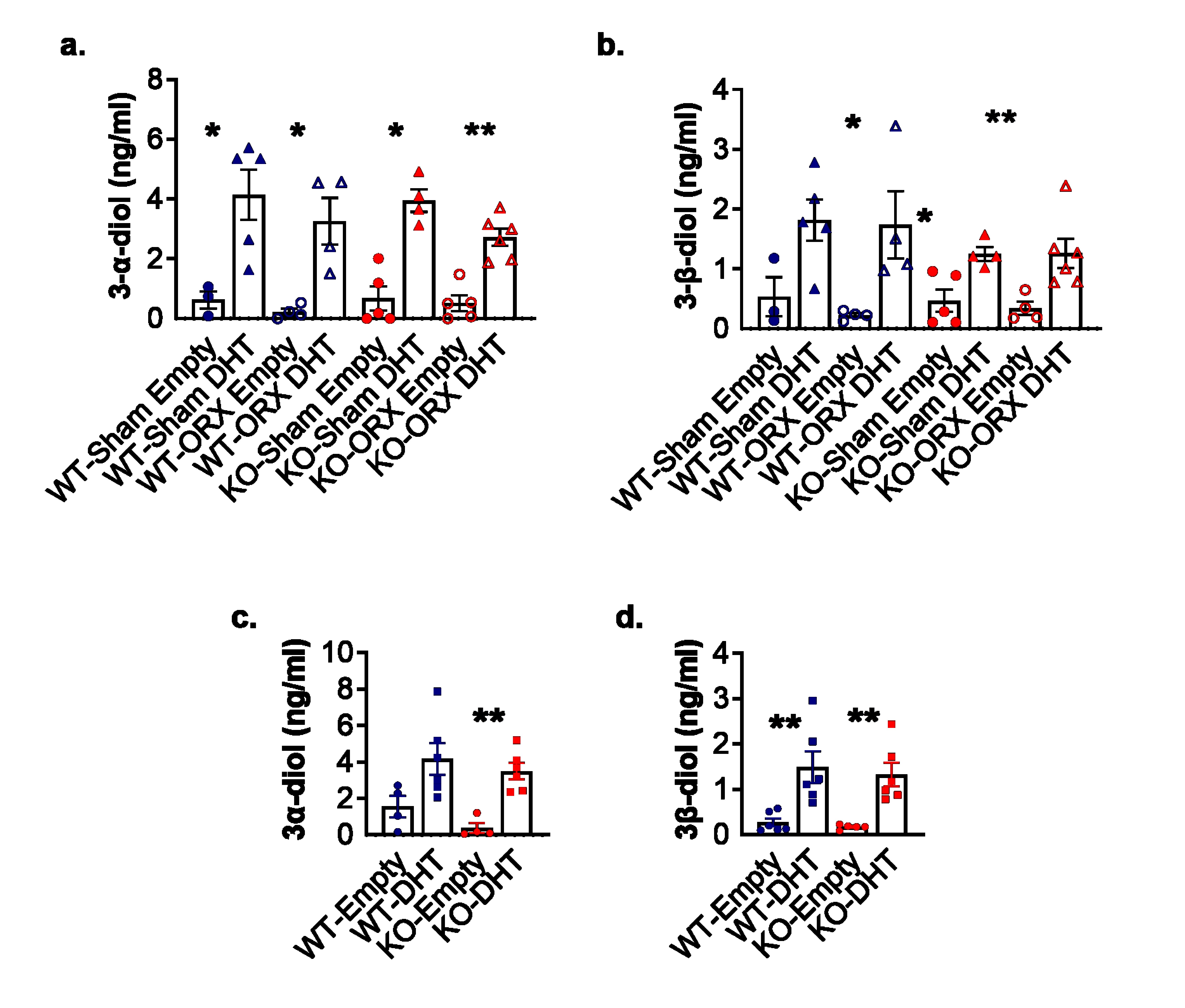

### Supplementary Fig 5

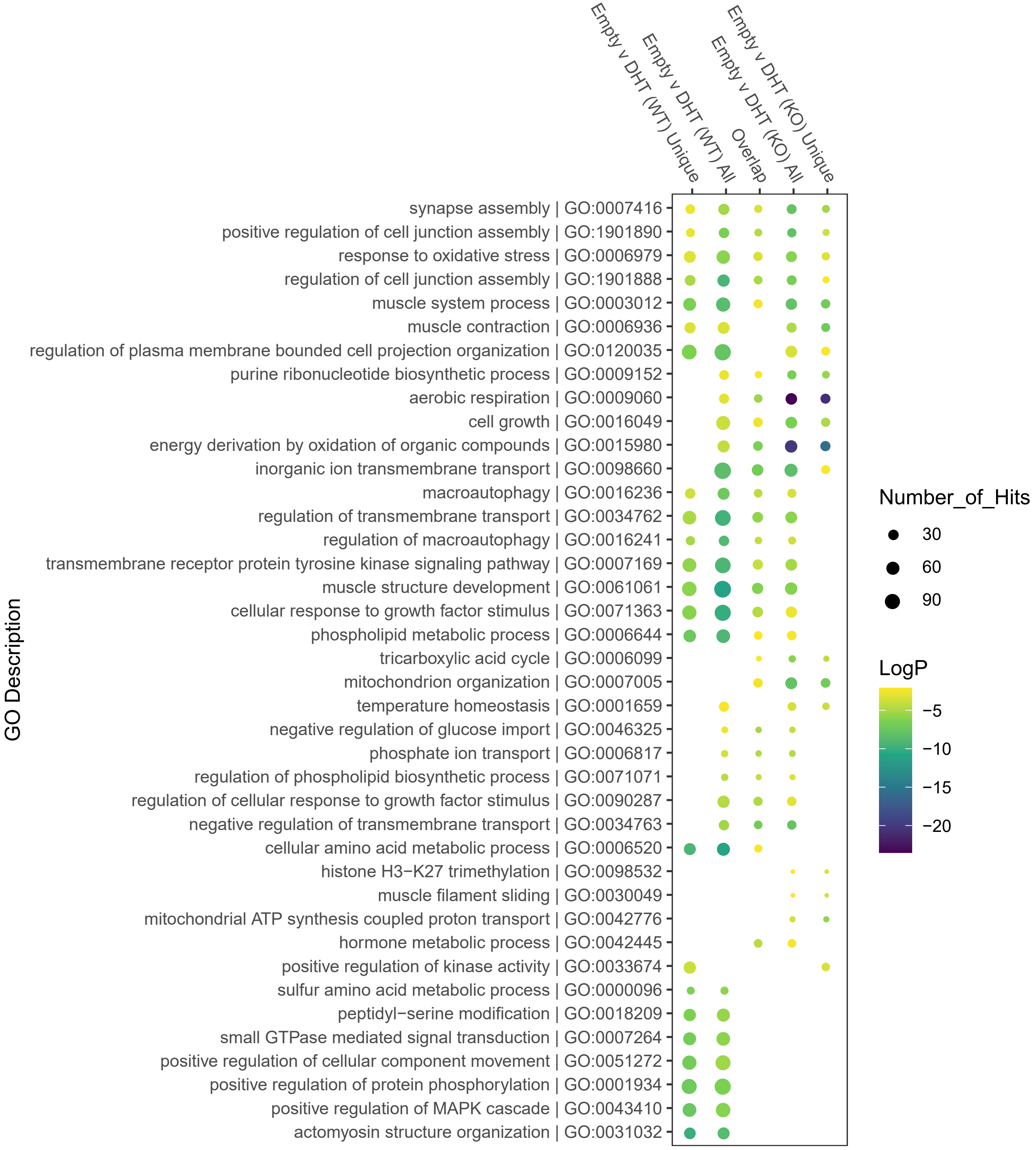
