## Supplementary Table for "*ACTN3* genotype influences androgen response in skeletal muscle"

**Supplementary Tables**

**Supplementary Table 1. Baseline expression of genes associated with Androgen, Estrogen and Thyroid receptor signalling.** *Ar, Smox, Odc1* are downregulated, but *Tceal7* is upregulated in *Actn3* KO relative to WT. In contrast, expression of genes associated with *Er* and *Tr* signalling are similar between WT and *Actn3* KO.

|  | **Gene** | **Gene name** | **Log fold change** | ***P* value** | **Adj P-value** |
| --- | --- | --- | --- | --- | --- |
| **Androgen signalling** | *Ar* | Androgen receptor | -0.29 | 0.0034 | 0.0252 |
|  | *Smox* | Spermine oxidase | -0.65 | 5.18E-6 | 1.99E-4 |
|  | *Odc1* | Ornithine decarboxylase 1 | -0.41 | 0.0010 | 0.0101 |
|  | *Tceal7* | Transcription Elongation Factor A Like | 2.32 | 5.81E-13 | 4.42E-10 |
| **Estrogen signalling** | *Esr1* | Estrogen receptor α | 0.09 | 0.4171 | 0.6165 |
|  | *Esrra* | Estrogen related receptor α | 0.10 | 0.1449 | 0.3169 |
|  | *Esrrb* | Estrogen related receptor β | -0.04 | 0.6659 | 0.8073 |
|  | *Esrrg* | Estrogen related receptor γ | 0.14 | 0.1415 | 0.3120 |
| **Thyroid receptor signalling** | *Thra* | Thyroid hormone receptor α | 0.07 | 0.1896 | 0.3765 |
|  | *Thrb* | Thyroid hormone receptor β | -0.02 | 0.8337 | 0.9145 |
|  | *Thrsp* | Thyroid hormone responsive | 0.39 | 0.0621 | 0.1828 |

Supplementary Table 2. DXA analysis of WT and *Actn3* KO body composition following 12 weeks of androgen deprivation

|  | WT | | | *Actn3* KO | | | Two-way ANOVA | | |
| --- | --- | --- | --- | --- | --- | --- | --- | --- | --- |
|  | SHAM  (*N*=9) | ORX (*N*=9) | *P*-value | SHAM (*N*=10) | ORX (*N*=8) | *P*-value | Genotype | Treatment | Interaction |
| Lean mass (g) | 23.28 ±2.17 | 20.03 ±2.20 | 0.0142 | 21.83±1.42† | 18.56 ±1.16 | <0.0001 | 0.017 | <0.0001 | 0.9900 |
| Fat mass (g) | 6.07 ±1.03 | 6.47 ±1.10 | 0.9494 | 6.48 ±1.10 | 7.71 ±2.23 | 0.2263 | 0.1430 | 0.1500 | 0.4590 |
| % Fat | 20.67 ±2.95 | 23.93 ±4.19 | 0.1084 | 22.81 ±3.48 | 28.9 ±5.25 | 0.0166 | 0.012 | 0.0010 | 0.2970 |
| BMD (g/cm^2^) | 0.053 ±0.0 | 0.049 ±0.0 | 0.0016 | 0.051 ±0.0* | 0.047 ±0.0 | <0.0001 | 0.0067 | <0.0001 | 0.7940 |
| BMC (g) | 0.409 ±0.02 | 0.362 ±0.04 | 0.0194 | 0.401 ±0.03 | 0.358 ±0.02 | 0.0031 | 0.5380 | <0.0001 | 0.8040 |

Supplementary Table 3. Effect of androgen deprivation on muscle mass and androgen responsive tissues.

|  | WT | | | *Actn3* KO | | | Two-way ANOVA | | |
| --- | --- | --- | --- | --- | --- | --- | --- | --- | --- |
|  | SHAM  (*N*=15) | ORX  (*N*=14) | *P*-value | SHAM  (*N*=19) | ORX  (*N*=15) | *P*-value | Genotype | Treatment | Interaction |
| SV (mg) | 375.99±39.16 | 25.51±18.85 | <0.0001 | 364.71±28.73 | 29.57±9.91 | <0.0001 | 0.5967 | <0.0001 | 0.2633 |
| LABC (mg) | 100.52±8.22 | 32.10±12.01 | <0.0001 | 109.28±7.11† | 30.62±10.77 | <0.0001 | 0.1372 | <0.0001 | 0.0363 |
| QUAD (mg) | 226.94±17.67 | 206.70±14.83 | 0.0037 | 193.25±10.7# | 164.34±12.49 | <0.0001 | <0.0001 | <0.0001 | 0.2252 |
| GST (mg) | 157.85±10.02 | 146.30±10.55 | 0.0059 | 140.1±5.76# | 116.7±7.33 | <0.0001 | <0.0001 | <0.0001 | 0.0075 |
| TA (mg) | 53.85±4.88 | 47.15±4.55 | 0.0008 | 50.47±3.13* | 41.91±4.03 | <0.0001 | 0.0002 | <0.0001 | 0.3892 |
| EDL (mg) | 11.08±0.86 | 9.88±1.52 | 0.0066 | 10.53±0.65 | 8.86±1.15 | <0.0001 | 0.0043 | <0.0001 | 0.3729 |
| SOL (mg) | 9.39±1.19 | 8.25±1.36 | 0.0826 | 9.92±0.65 | 8.61±1.07 | 0.8977 | 0.0002 | 0.1347 | 0.1693 |
| SPN (mg) | 223.87±24.04 | 180.82±13.01 | <0.0001 | 157.69±25.3# | 121.93±17.21 | <0.0001 | <0.0001 | <0.0001 | 0.5042 |
| Heart (mg) | 140.92±13.36 | 137.35±33.42 | 0.5556 | 153.32±29.09 | 124.21±19.71 | 0.2020 | 0.7612 | 0.8407 | 0.1134 |

Seminal vesicles (SV), levator ani bulbocavernosus (LABC), quadriceps (QUAD), gastrocnemius (GST), tibialis anterior (TA), extensor digitalis longus (EDL), soleus (SOL), spinalis (SPN) and heart (HRT). All samples were C57/BL6 male mice, had either SHAM or ORX surgery at aged 8-12 weeks. Data shown as mean ± SD. Mann-Whitney U pair-wise comparison tests.

**Supplementary Table 4.** Effect of *Actn3* genotype and DHT in male control and orchidectomised mice.

|  | WT | | | | | | *Actn3* KO | | | | | | Two-way ANOVA  (ORX +/- DHT) | | |
| --- | --- | --- | --- | --- | --- | --- | --- | --- | --- | --- | --- | --- | --- | --- | --- |
|  | SHAM+ Empty  (*N*=3) | SHAM  +DHT  (*N*=5) | *P*-value  SHAM +/- DHT | ORX+  Empty  (*N*=3) | ORX  +DHT  (*N*=4) | *P*-value  ORX +/- DHT | SHAM+  Empty  (*N*=4) | SHAM  +DHT  (*N*=4) | *P*-value  SHAM +/- DHT | ORX+  Empty  (*N*=5) | ORX  +DHT  (*N*=5) | *P*-value ORX +/- DHT | Geno-type | Treat-ment | Inter-action |
| BW (g) | 22.21  ±2.41 | 23.94  ±1.37 | 0.5714 | 19.58  ±0.21 | 25.40  ±3.08 | 0.0571 | 22.38  ±1.40 | 24.87  ±2.15 | 0.200 | 19.51 ±0.21 | 24.62  ±2.45 | 0.0079 | 0.6410 | 0.0002 | 0.8145 |
| SV (mg) | 237.23  ±2.90 | 353.10  ±0.37 | 0.0357 | 17.70  ±0.20 | 313.40  ±1.63 | 0.0571 | 232.28  ±0.56 | 365.30  ±0.50 | 0.0286 | 18.54  ±0.37 | 342.54  ±1.90 | 0.0286 | 0.4424 | <0.0001 | 0.5695 |
| LABC (mg) | 71.43  ±2.05 | 92.28  ±1.59 | 0.0357 | 14.4  ±3.93ᵟ | 100.95  ±3.83 | 0.0571 | 87.6  ±21.07 | 96.08  ±6.43 | 0.4857 | 15.78  ±4.76 | 103.82  ±6.46 | 0.0159 | 0.4121 | <0.0001 | 0.7711 |
| QUAD (mg) | 165.42 ±15.43 | 164.03  ±9.62 | 0.7857 | 147.58 ±12.70 | 179.23 ±21.39 | 0.1143 | 137.73  ±7.25 | 137.36  ±8.27 | 0.8857 | 123.16 ±6.59 | 140.89  ±12.66 | 0.8857 | 0.0005 | 0.0033 | 0.3310 |
| GST (mg) | 124.93  ±11.81 | 124.14  ±9.01 | >0.9999 | 113.82  ±6.45 | 129.61  ±11.38 | >0.9999 | 98.0  ±6.04 | 106.46  ±9.62 | 0.1143 | 90.79  ±4.70 | 108.84  ±2.11 | 0.1111 | <0.0001 | 0.0002 | 0.7386 |
| TA (mg) | 41.95 ±5.39 | 42.11  ±2.63 | >0.9999 | 35.05  ±0.95 | 42.88  ±4.22 | 0.0571 | 39.03  ±1.15 | 37.24  ±1.05 | 0.1143 | 33.66  ±1.03 | 40.31  ±3.42 | 0.0159 | 0.1867 | 0.0002 | 0.6855 |
| EDL (mg) | 9.38  ±2.41 | 8.53  ±1.37 | 0.7321 | 7.87  ±0.21 | 9.04  ±3.08 | 0.0571 | 8.58  ±1.40 | 8.54  ±2.12 | 0.6857 | 7.56  ±0.77 | 8.60  ±2.45 | 0.0635 | 0.2360 | 0.0525 | 0.8434 |
| SOL (mg) | 7.02  ±1.38 | 7.00  ±1.1 | >0.9999 | 6.28  ±0.33 | 7.40  ±1.03 | 0.7000 | 8.56  ±0.91 | 8.28  ±0.49 | 0.6857 | 7.24  ±0.87 | 9.03  ±0.64 | 0.1111 | 0.0066 | 0.0031 | 0.4096 |
| SPN (mg) | 158.60 ±18.08 | 174.70  ±16.47 | 0.3929 | 115.60  ±24.08 | 168.45  ±31.45 | 0.2286 | 117.26  ±18.62 | 117.65  ±13.36 | >0.9999 | 92.75  ±3.37 | 119.16  ±13.63 | 0.0159 | 0.0043 | 0.0023 | 0.2225 |
| HRT (mg) | 98.45 ±16.90 | 112.64  ±6.45 | 0.3810 | 96.00  ±4.76 | 121.88  ±4.01 | 0.0571 | 111.78  ±3.12 | 122.85  ±12.69 | 0.2286 | 92.92  ±3.66 | 117.76  ±13.07 | 0.0159 | 0.3806 | <0.0001 | 0.8981 |

ᵟWT-Sham Empty *vs.* WT-ORX Empty LABC (*P*=0.0159). Body weight (BW), seminal vesicles (SV), levator ani bulbocavernosus (LA), quadriceps (QUAD), gastrocnemius (GST), tibialis anterior (TA), extensor digitalis longus (EDL), soleus (SOL), spinalis (SPN) and heart (HRT). All samples were C57/BL6 male mice, had either Sham or ORX surgery and received an empty or DHT aged 4-5 weeks at time of implant. 3-5 animals per genotype/treatment. Data shown as mean ± SD. Mann-Whitney U pair-wise comparison tests.

**Supplementary Table 5.** Effect of *Actn3* genotype and DHT on body, muscle and heart mass in females.

|  | WT | | | *Actn3* KO | | | Two-way ANOVA | | |
| --- | --- | --- | --- | --- | --- | --- | --- | --- | --- |
|  | Empty  (*N*=6) | DHT  (*N*=6) | *P*-value | Empty  (*N*=6) | DHT  (*N*=6) | *P*-value | Genotype | Treatment | Interaction |
| BW (g) | 18.80 ±1.02 | 22.63 ±1.22 | 0.0022 | 19.51 ±1.47 | 20.48 ±2.29 | 0.4848 | 0.2759 | 0.0013 | 0.0382 |
| QUAD (mg) | 137. ±8.75 | 157.20 ±11.37 | 0.0087 | 124.60 ±12.65 | 127.70 ±13.52 | 0.8182 | 0.0003 | 0.0266 | 0.0962 |
| GST (mg) | 99.47±6.35 | 112.73±9.13 | 0.0087 | 86.32±9.13 | 91.73±8.49 | 0.3939 | <0.0001 | 0.0105 | 0.2497 |
| TA (mg) | 33.85 ±1.66 | 41.42±2.83 | 0.0022 | 34.18±3.27 | 35.42 ±3.18 | 0.5887 | 0.0227 | 0.0912 | 0.9874 |
| EDL (mg) | 7.11 ±1.02 | 8.49 ±1.22 | 0.0130 | 7.28 ±1.47 | 7.58 ±2.29 | 0.5887 | 0.2781 | 0.0187 | 0.1152 |
| SOL (mg) | 5.98 ±0.52 | 7.03 ±0.89 | 0.0216 | 6.81 ±0.59 | 6.85 ±0.75 | 0.7316 | 0.2651 | 0.0718 | 0.0944 |
| SPN (mg) | 124.65 ±7.31 | 181.3 ±30.78 | 0.0022 | 102.5 ±10.28^†^ | 105.75 ±15.17 | 0.6991 | <0.0001 | 0.0007 | 0.0020 |
| HRT (mg) | 89.03 ±5.90 | 112.02±5.60 | 0.0022 | 104.2±8.57^†^ | 105.13±12.03 | 0.9307 | 0.2543 | 0.0030 | 0.0055 |

†Baseline differences between WT and *Actn3* KO (Empty) (*P <* 0.05). Body weight (BW), quadriceps (QUAD), gastrocnemius (GST), tibialis anterior (TA), extensor digitalis longus (EDL), soleus (SOL), spinalis (SPN) and heart (HRT). All samples are from female C57/BL6 mice, aged 4-5 weeks at time of implant, 6 animals per genotype/treatment. Data shown as mean ± SD. Mann-Whitney U pair-wise comparison tests.

| ***Gene*** | **Protein** | **Description** | **Ref** |
| --- | --- | --- | --- |
| *Mybph* | Myosin binding protein H (MyBP-H) | MyBP-H is primarily expressed in fast-twitch muscle fibres and has a purported role in autophagy processes in cardiomyocytes as it colocalises with LC3 at the autophagosome membrane. Elevated expression of MyBP-H has also been associated with severe myopathy and muscle wasting in patients with amyotrophic lateral sclerosis and acute quadriplegic myopathy. | 1,2,3,4 |
| *Itpr1* | Inositol 1,4,5-triphosphate Receptor type 1 (IP_3_R1) | Knockdown of *Itpr1* inhibits C2C12 myoblast differentiation and impairs muscle regeneration in muscles of aged mice. Also a direct AR target gene in prostate cancer cells. | 5,6 |
| *Spns2* | Sphingosine-1 phosphate transporter spinster 2 | Increased expression of *Spns2* is paralleled by increases in atrogin-1 in dexamethasone-induced atrophy of C2C12 myotubes and in muscles of cachectic mice. | 7 |
| *Pitpna* | Phosphatidylinositol transfer protein-α | Integral for membrane trafficking and regulates phosphatidyl-inositol signalling between membrane compartments in eukaryotic cells. Repression of *Pitpna* has been shown to ameliorate the pathology of Duchenne muscular dystrophy. | 8,9 |
| *Syne1* | Nesprin-1 | Nesprin-1 is a key scaffolding protein involved in the mechanical coupling of the nucleus and the actin cytoskeleton and indirectly associates with various signalling pathways such as MAPK and calcineurin through its binding partners; mutations in nesprin-1 are involved in the pathogenesis of Emery Dreifuss muscular dystrophy. Also a direct AR target gene in prostate cancer cells. | 5,10,11 |
| *Ampd1* | AMP deaminase 1 | AMPD1 is essential for energy production through purine nucleotide interconversion and mutations cause metabolic myopathy in 2-3% of Caucasians. | 12 |
| *Ttll7* | Tubulin tyrosine ligase like-7 | Neurite growth | 13 |
| *2310016D23Rik* |  | ncRNA; function unknown | n/a |

**Supplementary Table 6.** Putative functions of differentially expressed genes that show significant interaction between *Actn3* genotype and androgen deprivation, as well as *Actn3* genotype and DHT. Many of these genes are also associated with muscle atrophy and disease.
